# Role of coiled-coil registry shifts in activation of human Bicaudal D2 for dynein recruitment upon cargo-binding

**DOI:** 10.1101/638536

**Authors:** Crystal R. Noell, Jia Ying Loh, Erik W. Debler, Kyle M. Loftus, Heying Cui, Blaine B. Russ, Puja Goyal, Sozanne R. Solmaz

## Abstract

Dynein adaptors such as Bicaudal D2 (BicD2) recognize cargo for dynein-dependent transport. BicD2-dependent transport pathways are important for brain and muscle development. Cargo-bound adaptors are required to activate dynein for processive transport, but the mechanism of action is elusive. Here, we report the structure of the cargo-binding domain of human BicD2 that forms a dimeric coiled-coil with homotypic registry, in which both helices are aligned. To investigate if BicD2 can switch to an asymmetric registry, where a portion of one helix is vertically shifted, we performed molecular dynamics simulations. Both registry types are stabilized by distinct conformations of F743. For the F743I variant, which increases dynein recruitment in the *Drosophila* homolog, and for the human R747C variant, which causes spinal muscular atrophy, spontaneous coiled-coil registry shifts are observed, which may cause the BicD2-hyperactivation phenotype and disease. We propose that a registry shift upon cargo-binding activates auto-inhibited BicD2 for dynein recruitment.

**Highlights:**

- Stable, bona fide BicD2 coiled-coils with distinct registries can be formed.
- We provide evidence that a human disease mutation causes a coiled-coil registry shift.
- A coiled-coil registry shift could relieve BicD2-autoinhibition upon cargo-binding.
- The ability to undergo registry shifts may be an inherent property of coiled-coils.

**In Brief:** Our results support that stable coiled-coils of BicD2 with distinct registries can be formed, and suggest a molecular mechanism for such registry switches. We provide evidence that disease-causing mutations in coiled-coils may alter the equilibrium between registry-shifted conformers, which we propose as a general mechanism of pathogenesis for coiled-coils.

**Graphical Abstract:** 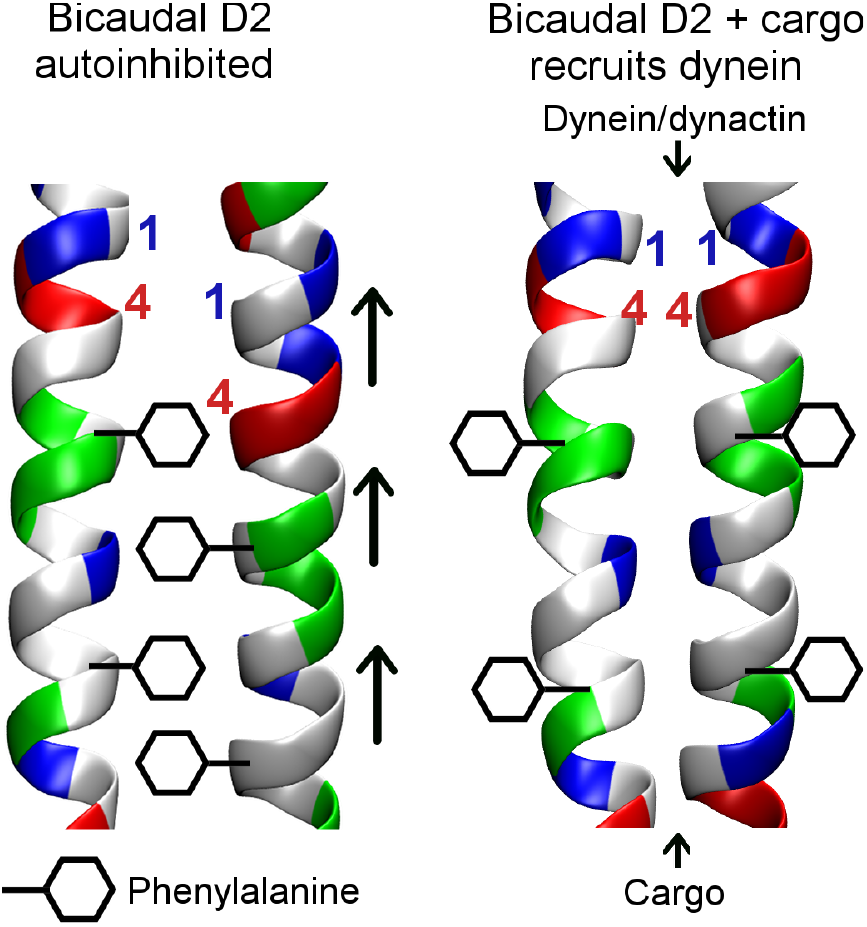

## INTRODUCTION

Cytoplasmic dynein is the predominant motor complex that mediates transport of almost all cargo that is transported towards the minus end of microtubules (Cianfrocco et al., 2015). Dynein adaptor proteins such as Bicaudal D2 (BicD2) (Hoogenraad et al., 2001) are an integral part of the dynein transport machinery, as they recognize cargo for transport and link it to the motor complex (Chowdhury et al., 2015; McKenney et al., 2014; Schlager et al., 2014a; Schlager et al., 2014b; Splinter et al., 2012; Urnavicius et al., 2015). Furthermore, in metazoans, cargo-loaded adaptor proteins such as BicD2 are required to activate dynein for processive transport, since they link dynein with its activator dynactin (Chowdhury et al., 2015; Urnavicius et al., 2015). In the absence of cargo, full-length BicD2 forms an auto-inhibited dimer (McKenney et al., 2014). In this conformation, the N-terminal domain (NTD) that contains a binding site for dynein and dynactin (Splinter et al., 2012) is inaccessible and likely masked by the C-terminal cargo binding domain (CTD) (Chowdhury et al., 2015; Liu et al., 2013; McClintock et al., 2018; McKenney et al., 2014; Schlager et al., 2014a; Schlager et al., 2014b; Sladewski et al., 2018; Splinter et al., 2012; Terawaki et al., 2015; Urnavicius et al., 2015). Once cargo binds to the CTD, auto-inhibition is released. This mechanism couples cargo loading with activation of transport (Chowdhury et al., 2015; McKenney et al., 2014; Schlager et al., 2014a; Schlager et al., 2014b; Splinter et al., 2012; Urnavicius et al., 2015). However, a structural framework for how dynein adaptors activate dynein for processive transport once cargo is bound has not been established. Of note, BicD2-autoinhibition is likely compromised by a classical dominant mutation of the *Drosophila* Bicaudal gene (F684I) that causes increased dynein recruitment to BicD2 (Liu et al., 2013; Mohler and Wieschaus, 1986). This variant results in anterior patterning defects, including double-abdomen embryos.

In humans, the predominant BicD2 cargoes are the GTPase Rab6^GTP^ (Matanis et al., 2002) and nuclear pore protein Nup358 (Splinter et al., 2010). Rab6^GTP^ recruits BicD2 to Golgi-derived and secretory vesicles. This transport pathway is important for signaling and neurotransmission at synapses (Matanis et al., 2002). Nup358 recruits BicD2 to the nucleus in G2 phase in order to position it respective to the centrosome during initial stages of mitotic spindle assembly. This pathway is essential for a fundamental process in brain development that is required for differentiation of certain brain progenitor cells (Baffet et al., 2015; Hu et al., 2013; Loftus et al., 2017; Splinter et al., 2010). The importance of BicD2-dependent transport events in brain and muscle development is reflected in the fact that human disease mutations cause a subset of cases of the neuromuscular disease spinal muscular atrophy (SMA), which is the most common genetic cause of death in infants (Martinez-Carrera and Wirth, 2015; Peeters et al., 2013; Synofzik et al., 2014). A structure of the human BicD2-CTD is currently not available but would be important to provide insights into underlying causes for SMA.

The structures of two homologs of the human BicD2-CTD (*Hs* BicD2) are currently known: mouse Bicaudal D1 (*Mm* BicD1) and *Drosophila* Bicaudal (*Dm* BicD) (Liu et al., 2013; Terawaki et al., 2015). In the structure of the *Mm* BicD1-CTD, equivalent knob residues in the two helices of the dimer are aligned at the same height, giving rise to a symmetric coiled-coil with a so-called homotypic registry (Terawaki et al., 2015). By contrast, the *Dm* BicD-CTD features an asymmetric coiled-coil structure, where one of the helices is shifted vertically with respect to the other one by one helical turn in the N-terminal region (heterotypic registry), whereas the helices are aligned in the C-terminal region (homotypic registry). It is unknown, whether the asymmetric coiled-coil registry is species-specific for *Dm* BicD or whether all BicD2 homologs undergo coiled-coil registry shifts in distinct functional states. Such a registry shift may play a role in the release of BicD2 auto-inhibition upon cargo binding.

Coiled-coil registry shifts have only been reported for a few structures (Carter et al., 2008; Croasdale et al., 2011; Gibbons et al., 2005; Macheboeuf et al., 2011; Snoberger et al., 2018; Xi et al., 2012), but may potentially be an inherent property of many coiled-coil structures. The most extensively studied example is dynein itself. In order to create the power stroke, distinct steps of the ATP hydrolysis reaction cycle in one domain must be coupled with microtubules binding and release in another domain. These two domains likely communicate through a coiled-coil registry shift in the connecting stalk domain (Carter et al., 2008; Gibbons et al., 2005). Molecular dynamics (MD) simulations are well-suited to investigate coiled-coil registry shifts, however, only one published study has applied this approach, but the resulting coiled-coil packing was not analyzed (Choi et al., 2011).

Here, we report the crystal structure of the human BicD2-CTD. Combined with our results from MD simulations of human BicD2 with homotypic and asymmetric registries, our data support the notion that BicD2 may undergo coiled-coil registry shifts. A registry shift upon cargo binding may activate BicD2 for dynein recruitment.

## RESULTS

### The structure of the human BicD2-CTD features a homotypic coiled-coil registry

In order to provide insights into underlying causes for spinal muscular atrophy (SMA) in human disease mutations of BicD2 (Peeters et al., 2013; Synofzik et al., 2014), we determined the x-ray structure of the human BicD2-CTD (residues 715-804) at 2.0 Å resolution (Figure 1). The R_Free-_factor is 25.7% (Table S1).

**Figure 1.**
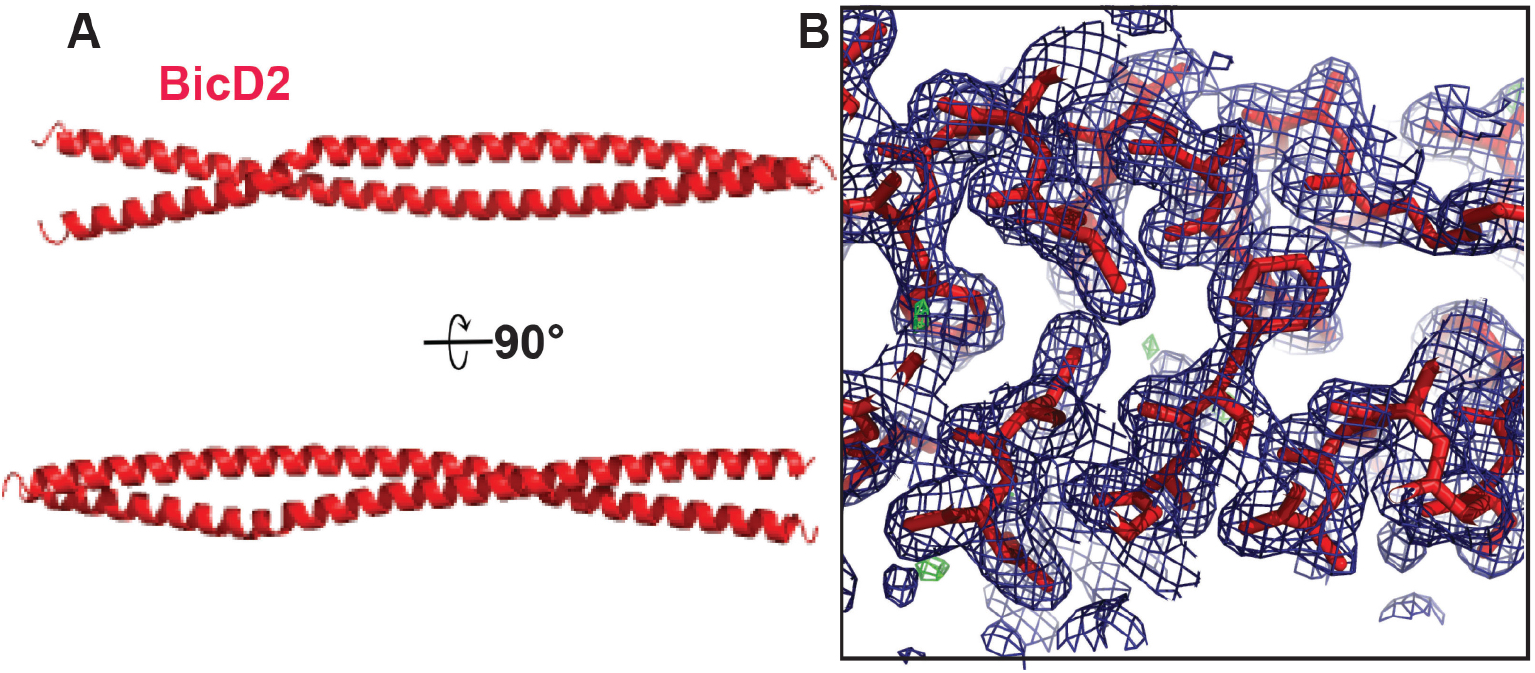
X-ray structure of human BicD2-CTD. (A) Structure of human BicD2-CTD in cartoon representation, rotated by 90°. (B) Structure of the BicD2-CTD in stick representation with 2f_O_-f_C_ electron density map (blue mesh). See also Table S1.

The BicD2-CTD forms a homo-dimeric coiled-coil in the crystal structure (Figure 1A) and also exists as a dimer in solution (Noell et al., 2018). Coiled-coils are characterized by heptad repeats ‘abcdefg’ in the protein sequence, where residues in the ‘a’ and ‘d’ positions of the heptad repeat are usually hydrophobic and engage in characteristic knobs-into-holes interactions at the core. Each knob residue fits into a hole composed of four residues on the opposite chain (Crick, 1953; O’Shea et al., 1991). The human BicD2-CTD features a homotypic coiled-coil registry, where identical knob residues at ‘a’ and ‘d’ positions of the heptad repeat from both chains intercalate to form a layer of knobs-into-holes interactions (Figure 2A, C, Figure S1A). In comparison, the *Dm* BicD-CTD coiled-coil has a homotypic registry in the C-terminal half (Figure 2D, green), where the α-helices are aligned, and a heterotypic registry in the N-terminal half, where each knob residue i from one chain intercalates with a knob residue i+4 in the second chain (Figure 2D, yellow) (Liu et al., 2013).

**Figure 2.**
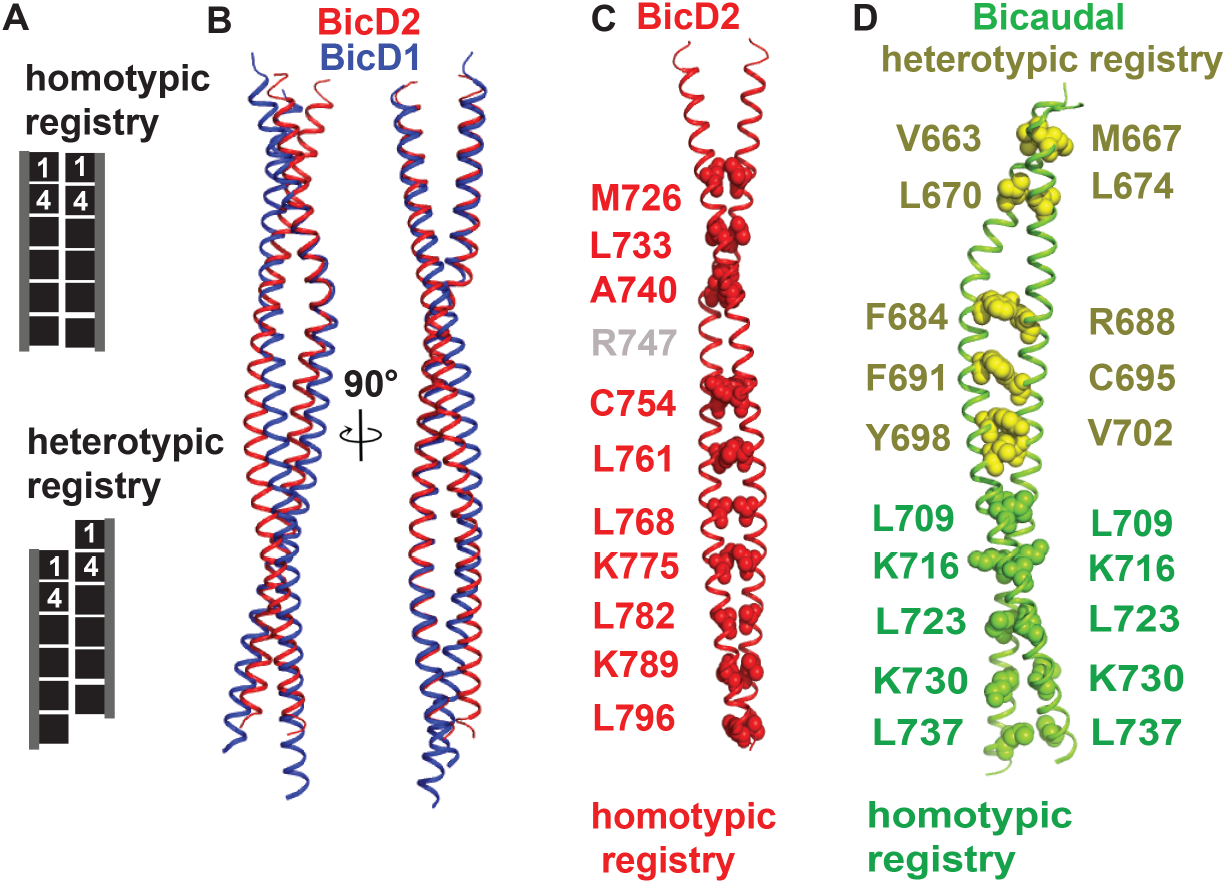
Human BicD2-CTD features a homotypic registry. (A) Schematic representation of homotypic and heterotypic coiled-coil registry. (B) Least squares superimposition of the structures of human BicD2-CTD (red) and mouse BicD1-CTD (blue, PDB ID 4YTD) (Terawaki et al., 2015), shown in cartoon representation. (C) Structure of human BicD2-CTD. Residues in ‘a’ position of the heptad repeat that form knobs-into-holes interactions are labeled in red and shown in spheres representation. (D) Structure of *Dm* BicD (PDB ID 4BL6) (Liu et al., 2013), which features a asymmetric registry (heterotypic registry in yellow, homotypic registry in green). See also Figure S1 and Table S2.

Interestingly, the structures of the human BicD2-CTD and the *Mm* BicD1-CTD (Terawaki et al., 2015), which have a homotypic registry (Figure S1), were obtained by co-crystallizing the proteins in the presence of cargo, whereas the *Dm* BicD-CTD with asymmetric registry was crystallized in absence of cargo. More specifically, our crystals of the human BicD2-CTD were obtained by crystallizing a minimal BicD2/Nup358 complex. Whether the presence of cargo during crystallization helps to stabilize a homotypic registry remains to be established. Despite the high homology of the *Hs* BicD2-CTD and *Mm* BicD1-CTD (90% sequence identity), the root mean square deviation (RMSD) for superimposition of the structures is 2.8 Å. This is similar to the RMSD for superimposition of *Hs* BicD2-CTD and *Dm* BicD-CTD (3.6 Å) (75% sequence identity). The RMSD indicates significant structural differences, which are observed especially at the dimer interface (Figure 2B, Table S2). The interface area is smaller for the *Dm* BicD-CTD (1764 Å^2^), compared to *Hs* BicD2 (2161 Å^2^) and *Mm* BicD1 (2176 Å^2^), which may make the *Dm* BicD-CTD less stable than its homologs. It should be noted that the smaller interface is likely due to the fact that the first few residues are not resolved in the structure of the *Dm* BicD-CTD.

Interestingly, in all BicD2 homologs the coiled-coil is interrupted between ‘a’ position residues A740 and C754 of BicD2 (Figure 2C, D, Figure S1), and thus residues in this region do not engage in knobs-into-holes interactions. This interruption likely serves to accommodate two bulky aromatic residues (F743, F750) at ‘d’ positions of the heptad repeat (Figure 3 A-E). Due to their large size, aromatic residues are rarely found at the core of coiled-coils. Residue R747 (‘a’-position) is sandwiched between F743 and F750. Y757 is located below, and engages in knobs-into-hole interactions (Figure 3 A-C). It was previously suggested that the asymmetric coiled-coil structure of *Dm* BicD is stabilized by specific interactions of F684, F691 and Y698 (Liu et al., 2013; Terawaki et al., 2015). This is supported by a comparison of the conformation of these residues in the three BicD2 homologs. In *Hs* BicD2, the aromatic rings of the F743 residues are stacked face-to-face, similar to the orientation observed in *Mm* BicD1 (Figure 3 B, C and Figure S1B). However, in *Dm* BicD, the aromatic rings of F684 from both chains are stacked edge-to-face (Figure 3 D, E). F750 and homologous residues form distinct conformations in the structures of all three homologs, allowing for structural plasticity (Figure 3, Figure S1B). Finally, knob residue Y757 forms similar interactions in *Mm* BicD1 and *Hs* BicD2, but the homologous residue Y698 has a different conformation in *Dm* BicD. This is in line with the suggestion that these three aromatic residues form distinct interactions which either stabilize the homotypic registry or the asymmetric coiled-coil registry (Liu et al., 2013; Terawaki et al., 2015).

Recently, a novel human disease mutation, R747C, was identified in the human BicD2 gene (Synofzik et al., 2014). Interestingly, R747 is located at the ‘a’ position of the heptad repeat in the area where the coiled-coil is interrupted (Figure 3 F, G). The positively charged arginine side chain of this residue is turned towards the surface of the coiled-coil, rather than engaging in knobs-into-holes interactions at the core (Figure 3 F, G). From the structural analysis, we predict that the R747C mutation destabilizes the structure of the coiled-coil. The smaller and more hydrophobic cysteine side chain may likely engage in knobs-into-holes interactions at the coiled-coil core, resulting in an interface that would not be able to accommodate the aromatic residues and destabilize the dimer.

**Figure 3.**
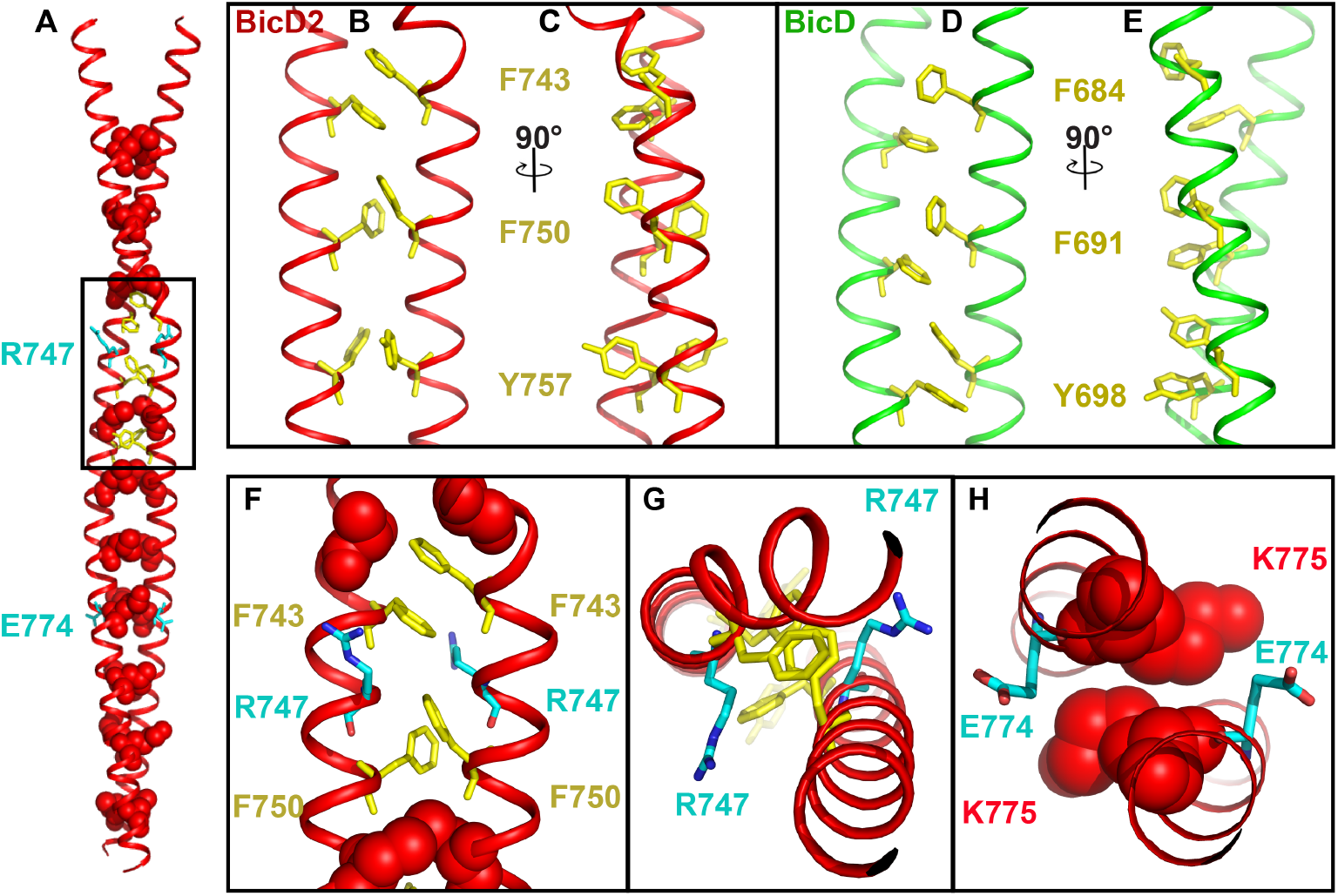
Role of aromatic residues in coiled-coil registry shift. (A) The BicD2-CTD is shown in the same representation as in Figure 2C. F743, F750 and Y757 are shown in yellow stick representation. R747 and E774, which are mutated in patients with SMA are shown in cyan stick representation and labeled. The boxed area is shown enlarged in (B, F) and rotated by 90° in (C). (D-E) *Dm* BicD-CTD (Liu et al., 2013) shown in the same representation, homologous aromatic residues are labeled. (F) Close-up view of human BicD2-CTD near R747. Top view is shown in (G). (H) Top view of E774 and nearby knob residue K775 (red sphere representation).

A second human BicD2 disease mutation is E774G (Peeters et al., 2013; Terawaki et al., 2015). The negatively charged glutamate side chain of E774 is oriented towards the surface (Figure 3H). Notably, E774 is close to residue K775, which is located at the ‘a’ position of the heptad repeat. A space-filling model of K775 illustrates that E774 is important to orient the lysine side chain correctly in order to form the knobs-into-holes interactions (Figure 3H). An E774G mutation is predicted to destabilize the structure of the BicD2 coiled-coil, as glycine is a very flexible residue that has a somewhat destabilizing effect when incorporated in α-helices.

To conclude, the BicD2-CTD has a homotypic coiled-coil registry, which is stabilized by distinct conformations of aromatic residues. Furthermore, our structural analysis suggests that the human disease mutations R747C and E774G that cause SMA destabilize the coiled-coil structure of BicD2.

### In MD simulations of BicD2, spontaneous coiled-coil registry shifts are observed for the human disease variant R747C and for the classical *Drosophila* Bicaudal allele

In order to compare the relative stability of conformations of human BicD2 with distinct coiled-coil registries (homotypic registry, asymmetric registry, and heterotypic registry), computational studies were carried out. A simulation of the homotypic coiled-coil registry starting from the crystal structure of the human BicD2-CTD maintained a homotypic coiled-coil registry throughout the simulation, with the knob residues at ‘a’ and ‘d’ positions of the heptad repeat depicted in Figure 4A. Analysis of the equilibrated structure revealed a key difference in the conformation of F743 compared to the starting crystal structure. The F743 side chains from both monomers that interact within the hydrophobic core of the dimer in the crystal structure rotated out to be solvent-exposed within ~0.3 ns of the simulation, and adopted a conformation similar to that of F750 (Figure 4C). This led to changes of the helical curvature compared to the crystal structure.

To investigate whether human BicD2 could assume a conformation with an asymmetric (homotypic/heterotypic) registry, this conformation was created *in silico* by mutating the x-ray structure of the *Dm* BicD-CTD (Liu et al., 2013) to human BicD2. In addition, we created a model for human BicD2 with heterotypic registry by pulling one helix of the dimer downwards respective to the other by one helical turn. The resulting structures were equilibrated using MD simulations. The registries remained stable in the simulations and the equilibrated structures exhibited a comparable number of knobs-into-holes interactions as the experimentally determined structures, with the knob residues at the ‘a’ and ‘d’ positions of the heptad repeat depicted in Figure 4B and Figure S2. In conjunction with excellent structure validation results, and an absence of steric clashes (Table S3), this confirms that the resulting BicD2 structures with the asymmetric and heterotypic registries are *bona fide* coiled-coils (Figure S3).

**Figure 4.**
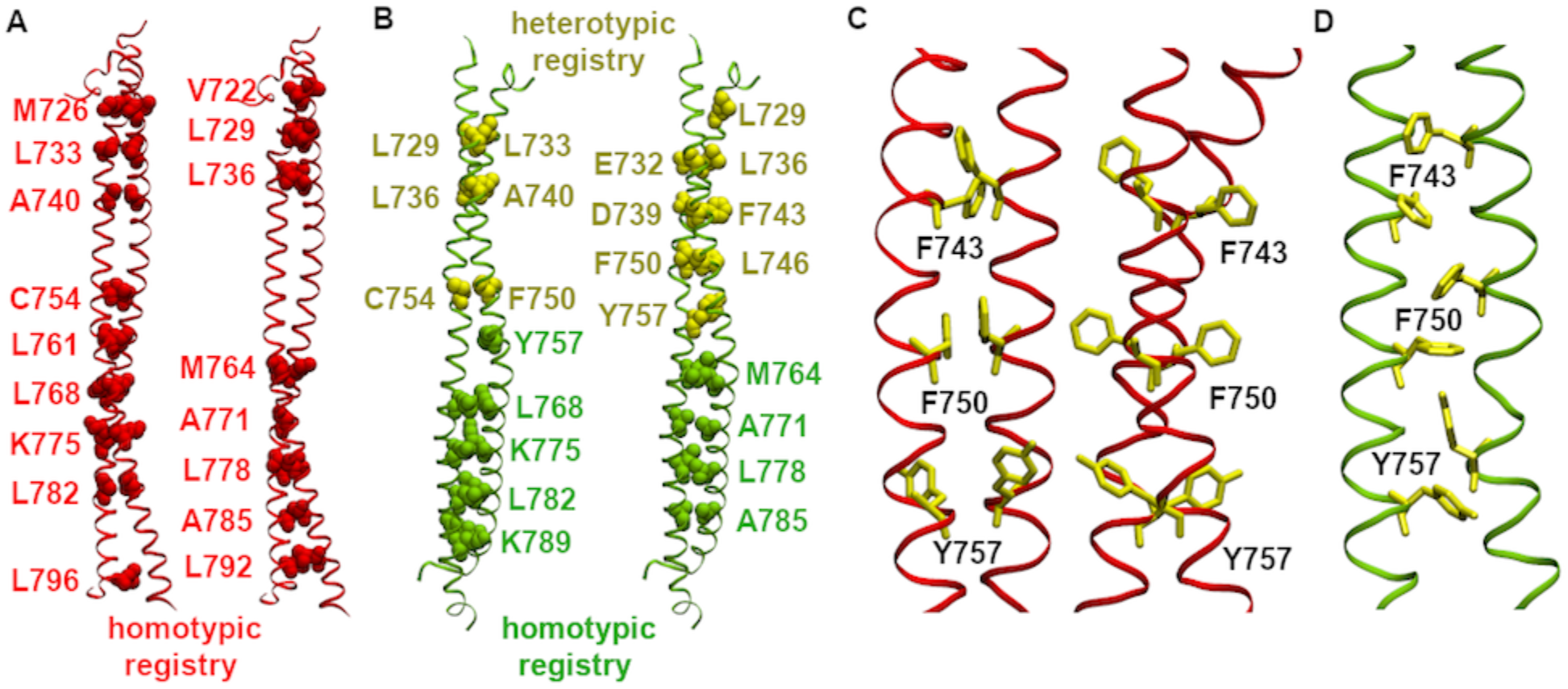
Asymmetric registry for human BicD2-CTD created *in silico* has a coiled-coil structure, and the registry type is correlated with the orientation of key aromatic residues. (A-D) Equilibrated structures of human BicD2-CTD with homotypic (A, C) and asymmetric (B, D) registries in cartoon representation at 300 K. (A, B) ‘a’ and ‘d’ position residues are shown as spheres in the left and right panel, respectively. (A) Homotypic registry. (B) Asymmetric registry (heterotypic in yellow, homotypic in green). (C, D) The key aromatic residues are depicted using a stick model. (C) Note that F743, which is part of the dimer interface in the crystal structure with homotypic registry (Figure 3) rotates outwards in ~0.3 ns of simulation. F750 and Y757 point outwards into solution as observed in the crystal structure. (D) In the structure with asymmetric registry, F743, F750 and Y757 are all part of the dimer interface. See also Figure S2, Figure S3 and Table S3.

**Table 1.** Dimer interfaces for BicD2-CTD with homotypic and asymmetric coiled-coil registries^a^. See also Table S4.

| BicD2-CTD, homotypic |  |  | BicD2-CTD, asymmetric |  |  |
| --- | --- | --- | --- | --- | --- |
| Chain A | Chain B | Dist. (Å) | Chain A | Chain D | Dist. (Å) |
| **Hydrogen bonds** |  |  | **Hydrogen bonds** |  |  |
| SER 712 | LYS 719 | 3.57 | GLU 724 | GLU 718 | 1.94 |
| OG | O |  | H | OE2 |  |
| ARG 795 | GLU 800 | 1.86 | ARG 730 | THR 725 | 1.91 |
| HE | OE2 |  | HH11 | OG1 |  |
| ARG 795 | GLU 800 | 1.77 | LYS 737 | GLU 732 | 1.80 |
| HH21 | OE1 |  | HZ3 | OE1 |  |
| GLU 774 | LYS 775 | 1.78 | ARG 747 | ASP 739 | 1.80 |
| OE2 | HZ3 |  | HH11 | OD1 |  |
| GLN 788 | LYS 789 | 1.94 | LYS 775 | GLU 774 | 1.80 |
| OE1 | HZ3 |  | HZ2 | OE1 |  |
| GLU 797 | ARG 795 | 1.89 | LYS 789 | GLN 788 | 1.97 |
| OE1 | HE |  | HZ1 | OE1 |  |
| GLU 797 | ARG 795 | 2.02 | ARG 795 | LEU 799 | 1.81 |
| OE1 | HH21 |  | HE | O |  |
|  |  |  | ARG 795 | LEU 799 | 2.01 |
|  |  |  | HH21 | O |  |
|  |  |  | ASP 755 | ARG 753 | 1.80 |
|  |  |  | OD1 | HH12 |  |
|  |  |  | CYS 754 | ARG 753 | 1.92 |
|  |  |  | O | HH11 |  |
|  |  |  | GLU 774 | LYS 775 | 1.83 |
|  |  |  | OE2 | HZ3 |  |
|  |  |  | GLN 788 | LYS 789 | 1.82 |
|  |  |  | OE1 | HZ1 |  |
|  |  |  | LEU 799 | ARG 795 | 2.14 |
|  |  |  | O | HE |  |
|  |  |  | LEU 799 | ARG 795 | 1.96 |
|  |  |  | O | HH21 |  |
| **Salt bridges** |  |  | **Salt bridges** |  |  |
| ARG 795 | GLU 800 | 3.61 | LYS 737 | GLU 732 | 2.80 |
| NE | OE1 |  | NZ | OE1 |  |
| ARG 795 | GLU 800 | 2.88 | ARG 747 | ASP 739 | 2.78 |
| NE | OE2 |  | NH1 | OD1 |  |
| ARG 795 | GLU 800 | 2.79 | LYS 775 | GLU 774 | 2.82 |
| NH2 | OE1 |  | NZ | OE1 |  |
| ARG 795 | GLU 800 | 3.57 | ARG 795 | LEU 799 | 2.79 |
| NH2 | OE2 |  | NE | O |  |
| GLU 774 | LYS 775 | 2.77 | ARG 795 | LEU 799 | 2.91 |
| OE2 | NZ |  | NH2 | O |  |
| GLU 797 | ARG 795 | 2.85 | ASP 755 | ARG 753 | 2.78 |
| OE1 | NE |  | OD1 | NH1 |  |
| GLU 797 | ARG 795 | 2.92 | ASP 755 | ARG 753 | 3.94 |
| OE1 | NH2 |  | OD1 | NH2 |  |
|  |  |  | GLU 774 | LYS 775 | 2.76 |
|  |  |  | OE2 | NZ |  |
| **Interface Area (Å<sup>2</sup>)** |  |  | **Interface Area (Å<sup>2</sup>)** |  |  |
| 2302 |  |  | 2477 |  |  |

<sup>a</sup> Structures were energy-minimized post equilibration at 300 K. The structure with the asymmetric registry has 8 and 9 fewer residues in chain A and B, respectively, compared to the structure with homotypic registry, due to varying residue numbers in the original crystal structures. Similar analysis for the structure with heterotypic registry is provided in Table S4.

In order to investigate which conformation of human BicD2 is the most stable, we analyzed the dimer interface areas of the equilibrated structures with different registries (Table 1 and Table S4). As the interface is mainly stabilized by hydrophobic interactions, the interface area can be an indicator of overall dimer stability. The asymmetric registry has the largest interface area (2477 Å^2^), followed by the heterotypic (2339 Å^2^) and the homotypic (2302 Å^2^) registry. The larger interface area of the asymmetric registry compared to the homotypic registry can be understood partly by the difference in the orientation of the F743, F750, and Y757 side chains. In terms of other non-covalent interactions, the asymmetric registry has the largest number of salt bridges and hydrogen bonds (8/14 respectively), followed by the homotypic (7/7) and the heterotypic (5/7) registry. However, even though the asymmetric registry is likely the most stable conformation, overall, the difference compared to the homotypic registry is rather small, suggesting that the free energy of both conformations is not substantially different.

**Figure 5.**
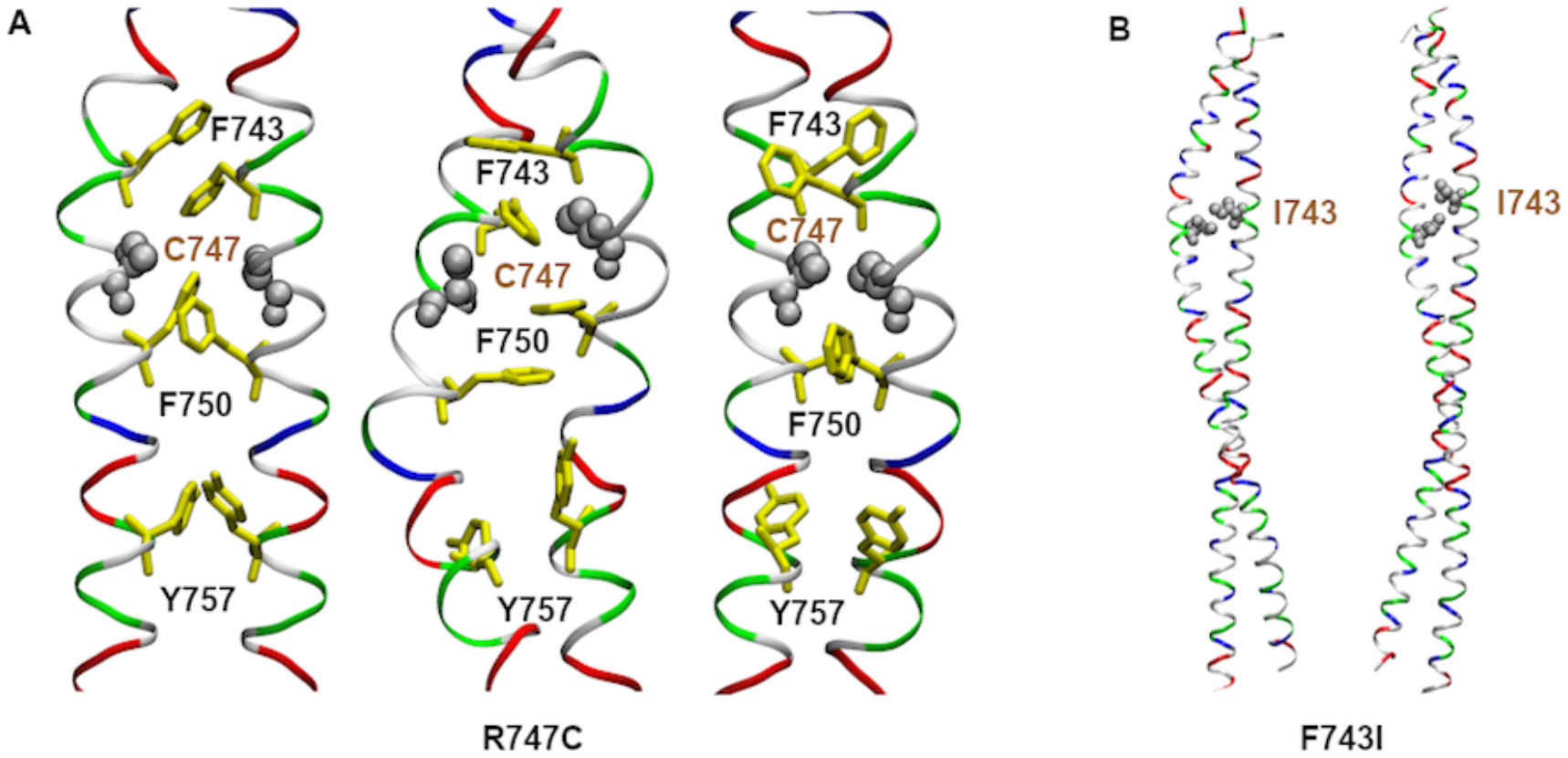
R747C and F743I mutations cause short-lived local and long-lived global registry shifts, respectively, in MD simulations of human BicD2-CTD. (A) Left: Cartoon representation of the equilibrated human BicD2-CTD structure with homotypic registry, colored by residue type (blue: positively charged, red: negatively charged, green: polar, white: non-polar). R747 was mutated to cysteine (silver spheres). F743 is part of the dimer interface, while F750 and Y757 point outwards into solution. Middle: After ~30 ns of simulation, a local registry shift to heterotypic registry is induced, and F750 becomes part of the dimer interface. Right: After ~79 ns of simulation, F743 and F750 point outwards into solution and the registry returns to homotypic. (B) Left: F743I mutation (silver spheres) in the equilibrated structure of human BicD2-CTD with asymmetric (heterotypic/homotypic) registry. Right: The fully heterotypic coiled-coil registry after ~92 ns of simulation is shown. See also Table S4 and Figure S3.

A notable difference between BicD2 conformations with different registries is that each registry is stabilized by distinct side-chain orientations of F743 and F750. Of note, in the equilibrated homotypic registry, the F743 and F750 side chains are rotated out to be solvent-exposed (Figure 4C). In contrast, in the asymmetric (Figure 4D) and heterotypic (Figure S2) registries, these side chains are rotated towards the dimer core with a stabilizing edge-to-face interaction between the two F743 side chains and the two F750 side chains. The Y757 side chains also adopt different orientations in the three registries. While their hydroxyl groups are oriented towards the solvent in the homotypic and heterotypic registries, these groups roughly point in the same direction in the asymmetric registry. These differences suggest that F743 and F750 may be key residues involved in a registry shift, as proposed earlier (Liu et al., 2013).

This hypothesis was confirmed by simulations of human BicD2 with the R747C mutation, which causes spinal muscular atrophy (Synofzik et al., 2014). R747 is sandwiched between residues F743 and F750. During simulation of the R747C mutant, a spontaneous local coiled-coil registry shift from homotypic to heterotypic was observed, with recovery of the homotypic registry later in the simulation. For the duration of ~79 ns, the F743 side chain in either monomer remained oriented towards the hydrophobic core of the dimer and a local registry shift around F743 from homotypic to heterotypic was induced. This was accompanied by an inward rotation of the F750 side chains. Subsequently, the F743 and F750 side chains rotated outwards and became exposed to the solvent, which led to recovery of the homotypic registry in this region (Figure 5A, Figure S3). These results suggest that distinct orientations of the F743 and F750 side chains are specific to BicD2 with homotypic and heterotypic registry, at least locally. Furthermore, the fact that a spontaneous local coiled-coil registry shift is observed during simulation of the R747C variant supports the idea that human BicD2 is capable of coiled-coil registry shifts.

Simulations of the F743I mutant provided further support for this idea. The analogous F684I mutation in *Drosophila* is known to lead to increased association of *Dm* BicD with dynein, thereby “hyperactivating” BicD2-dependent transport (Liu et al., 2013). In order to gain insight into the effect of this mutation on the structure of human BicD2, an MD simulation for the F743I mutant of human BicD2 was carried out. Within ~92 ns of the simulation, the starting asymmetric homotypic-heterotypic registry changed to a fully heterotypic registry (Figure 5B, Figure S3, Table S4). This result indicates that the registry of the coiled-coil may switch in the F684I mutant, which may be the cause of the observed stronger association of *Dm* BicD with dynein.

### Human disease mutations lead to misfolding of BicD2

Our structural analysis suggests that the BicD2-CTD and the *Dm* BicD-CTD vary in their stability and that the human disease mutations E774G and R747C destabilize the coiled-coil structure. To test these hypotheses, we characterized these proteins by circular dichroism (CD) spectroscopy to compare the thermodynamic stability of these proteins.

CD wavelength spectra of purified BicD2-CTD and *Dm* BicD-CTD are very similar and show minima at 208 and 222 nm, which are characteristic of α-helical proteins, which disappear upon denaturation at 95°C (Figure 6A). Furthermore, we probed the stability of the BicD2-CTD and *Dm* BicD-CTD by melting curves, where unfolding was monitored by CD spectroscopy at 222 nm and apparent melting temperatures (T_M_) were determined. The T_M_ for BicD2-CTD is 46.0±1.8°C, whereas the T_M_ of *Dm* BicD is 41.2±0.6°C (Figure 6B). Notably, the small difference of the melting temperature suggests that the stabilizing free energy of these two states is similar. Interestingly, the melting curve of the *Dm* BicD-CTD has a much steeper slope compared to the BicD2-CTD, suggesting distinct unfolding kinetics. Furthermore, the α-helical content of the BicD2-CTD decreases between temperatures of 4 – 25°C by 10%. These structural changes are fully reversible. CD-wavelength scans of BicD2-CTD recorded at 4°C are undistinguishable before and after the sample is subjected to a temperature gradient from 4 - 25°C (Figure 6C). These data support that the structure of the BicD2-CTD is dynamic and can undergo conformational changes in solution.

Two mutations of the human BicD2 gene were described that cause SMA and are located in the BicD2-CTD (R747C, and E774G) (Peeters et al., 2013; Synofzik et al., 2014). Our structural analysis suggests that these mutations destabilize the coiled-coil structure. Of note, purification of the BicD2-CTD/R747C and E774G variants resulted in a lower yield compared to the wild-type (Figure S4A), which is in line with lower stability. Furthermore, gel filtration profiles of the E774G variant indicated significant formation of high-molecular weight aggregates (data not shown). CD wavelength scans of the E774G disease variant lack the well-defined minima at 208 and 222 nm of α-helical proteins altogether, indicating misfolding (Figure 6D).

**Figure 6.**
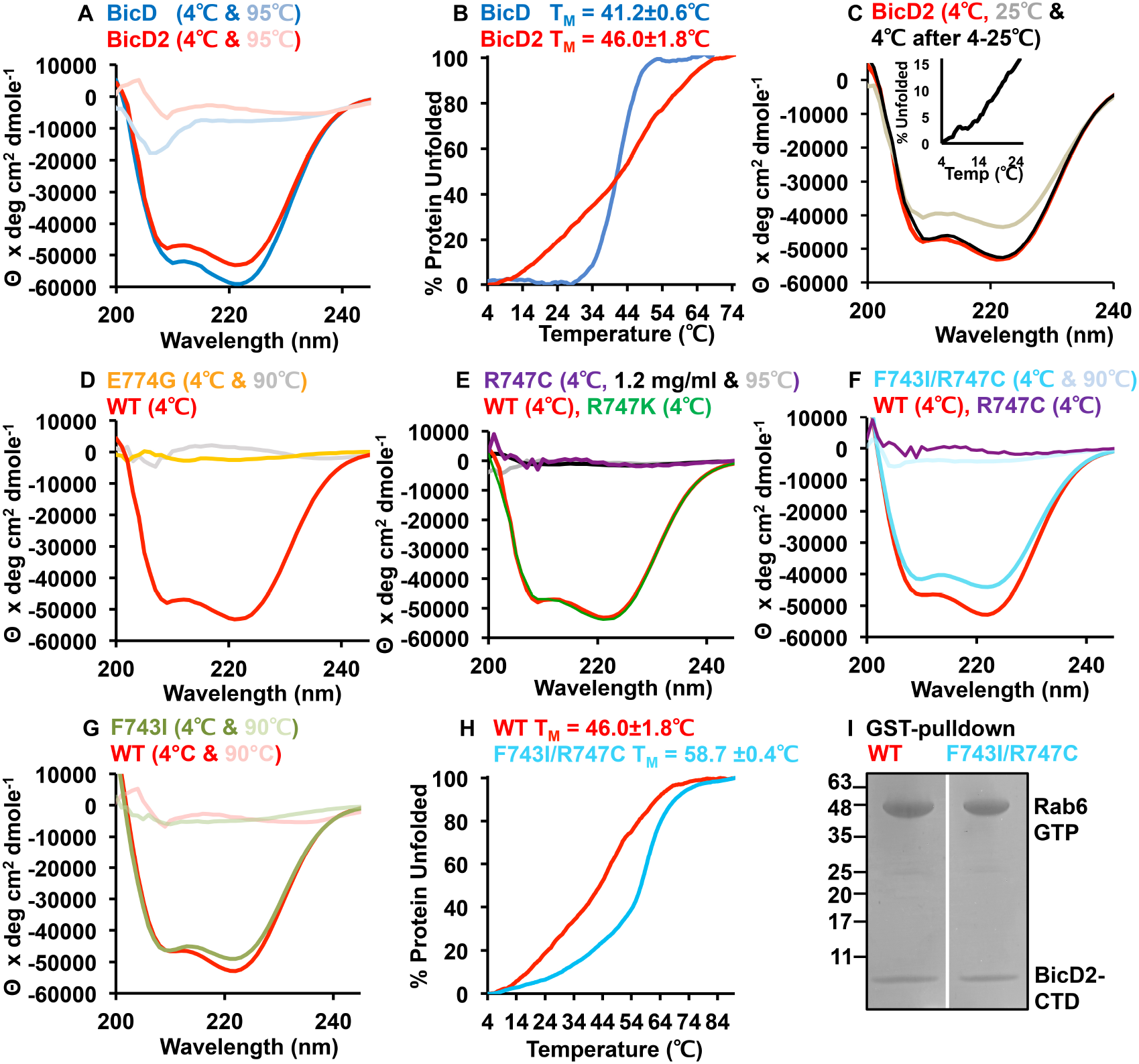
Human disease variants of BicD2 are misfolded. (A) CD wavelength scans of human BicD2-CTD (red) and *Dm* BicD-CTD (blue) at 4°C (native) and 95°C (random coil). The mean residue molar ellipticity ⊝ versus the wavelength is shown. (B) Thermal unfolding curves of BicD2-CTD (red), and *Dm* BicD (blue) monitored by CD spectroscopy at 222 nm. 0% and 100% Protein unfolded represents the minimum and maximum value of ⊝, respectively. Apparent melting temperatures T_M_ are shown. (C) The BicD2-CTD was subjected to a temperature gradient from 4 - 25°C (top insert), and cooled back to 4°C. CD wavelength spectra at 25°C (grey) and at 4°C before (red) and after the experiment (black) are shown. (D-H) CD wavelength spectra of wild-type BicD2-CTD (red) and its variants are shown. (D) BicD2-CTD/E774G. (E) BicD2-CTD/R747C (0.3 mg/ml at 4°C in purple, 0.6 mg/ml at 4°C in black, 0.3 mg/ml at 95°C in gray). The spectra are overlaid with the conservative R747K variant (green). (F) BicD2-CTD/F743I/R747C (overlaid with WT and R747C variant). (G) BicD2-CTD/F743I. (H) Thermal unfolding curves of BicD2-CTD/F743I/R747C (cyan) and WT (red), monitored by CD spectroscopy at 222 nm. (I) To assess binding, a GST-pulldown assay was performed with GST-tagged Rab6^GTP^ and BicD2-CTD, as well as its F743I/R747C variant. SDS-PAGE analysis of the elution fraction is shown, the location of molar mass standards (kDa) is indicated. See also Figure S4.

Notably, CD wavelength scans of the BicD2/R747C disease variant lack the characteristic α-helical minima at 208 and 222 nm, suggesting that this variant also leads to misfolding (Figure 6E). As a control, we also purified a conservative R747K variant. CD wavelength spectra and melting curves of the conservative R747K variant are very similar to the wild-type (TM=47.4±1.8°C and TM=46.0±1.8°C; Figure 6E, Figure S4B).

These results suggest that the BicD2-CTD/E774G and R747C variants are to a large degree misfolded to a random coil and confirm that these mutations destabilize the coiled-coil structure, which may be an underlying cause for SMA in these variants.

It should be noted that in our MD simulations of the BicD2-CTD/R747C variant, a spontaneous and transient registry shift is observed (Figure 5). Such transitions between registry-shifted conformers may destabilize BicD2, however, it is conceivable that misfolding occurs on a slow time scale that cannot be observed during MD simulations. In order to stabilize the R747C disease variant, we combined this registry-shift inducing mutation with a second mutation, F743I. Since the F743I mutation induces a spontaneous, but stable registry shift in BicD2-CTD during our MD simulations (Figure 5), we hypothesized that the resulting F743I/R747C double mutant could rescue folding and stabilize BicD2 in a registry-shifted conformer. Notably, CD wavelength spectra of the BicD2-CTD/F743I/R747C double mutant overlay well with the wild type, suggesting comparable α-helical content and confirming that the combination of these two variants rescues folding of BicD2 (Figure 6 F). As a control, we recorded CD wavelength spectra of the BicD2-CTD/F743I variant (Figure 6 G), which are comparable to both the wild type and the BicD2-CTD/F743I/R747C double variant spectra. Finally, we determined the apparent melting temperature T_M_ of the BicD2-CTD/F743I/R747C double variant (Figure 6 H). Its T_M_ indicates that this double variant is substantially more stable (T_M_= 58.5±0.5°C) compared to the wild type (T_M_=46.0±1.8°C). The double variant has also a higher T_M_ compared to the F743I single variant (T_M_=51.2±0.8°C, Figure S4). This stabilization effect of the double variant is not due to disulfide bond formation, since the BicD2 sample is kept in a reduced state. Next, we assessed effects of the F743I/R747C double mutation on binding of cargoes. We performed a pulldown assay with GST-tagged Rab6^GTP^ and the BicD2-CTD WT, as well as its F743I/R747C variant. Notably, a comparable amount of the wild type and the F743I/R747C variant is pulled down by Rab6^GTP^, indicating that the F743I/R747C mutation does not affect binding of cargoes (Figure 6 I).

To conclude, the BicD-CTD/R747C disease variant, which causes spinal muscular atrophy, is to a large degree misfolded. Our results confirm that the combination of the R747C mutation with the F743I mutation that also induces a registry shift in MD simulations of BicD2 (Figure 5), rescues folding of BicD2 and even stabilizes it substantially compared to the wild type (Figure 6). Notably, the interaction with the cargo Rab6^GTP^ is not altered by the F743I/R747C double mutation compared to the wild type.

## DISCUSSION

We report here the x-ray structure of the cargo-binding domain of human BicD2, which provides insights into the molecular basis of spinal muscular atrophy (SMA) in the human disease variants R747C and E774G. Our results suggest that these mutations destabilize the structure of the coiled-coil and lead to misfolding, which remains to be confirmed in the context of the full-length protein. The misfolding likely results in loss of function, which is further supported by previous reports of cargo-binding defects (Peeters et al., 2013; Synofzik et al., 2014; Terawaki et al., 2015). Our data emphasize the fact that these mutations tamper with the coiled-coil structure rather than altering cargo binding specifically.

To investigate if BicD2 could switch from a homotypic registry, where equivalent knob residues from both α-helices are aligned (as observed in our crystal structure), to an asymmetric registry, in which a portion of one helix slides vertically against the other, we performed MD simulations. In these simulations, both types of registries are stabilized by distinct conformations of residues F743 and F750. Interestingly, for the human disease variant R747C, a spontaneous local registry shift was observed, which is facilitated by specific conformations of F743. Human disease mutations have so far not been implicated in causing registry shifts in coiled-coil structures; however, our data suggest that registry shifts should indeed be considered as a general disease mechanism for this class of proteins.

While we observed misfolding of the BicD2-CTD/R747C variant upon recombinant protein expression, a spontaneous and transient registry shift is observed for this R747C variant in our MD simulations. Such transitions between registry-shifted conformers may destabilize the structure of BicD2 and ultimately lead to misfolding. Misfolding was not observed in our MD simulations, likely because it may occur on a slow time scale that cannot be covered by these simulations. In order to stabilize the R747C disease variant, we combined it with the F743I mutation, which induces a spontaneous, but stable registry shift in MD simulations. We hypothesized that the resulting F743I/R747C double variant could rescue folding and stabilize BicD2 in a registry-shifted conformer, which was indeed supported by our results. The F743I/R747C double variant even interacts with its cargo Rab6^GTP^ at comparable strength as observed for the wild type protein, further confirming that the R747C mutation does not interfere with cargo binding.

Notably, cargo-bound adaptors such as BicD2 are required to activate dynein for processive transport. In order to prevent non-productive transport events in the absence of cargo, BicD2 exists in an auto-inhibited form. A registry shift in BicD2 upon cargo binding could propagate through the coiled-coil to the N-terminal dynein-binding site and is a plausible mechanism to relieve autoinhibition of BicD2 in order to activate it for dynein recruitment. Our data support the notion that human BicD2 may undergo coiled-coil registry shifts. BicD2 conformations with homotypic and asymmetric registries are stable in MD simulations. Furthermore, these structures feature good stereochemistry, no steric clashes, as well as characteristic knobs-into-holes interactions, confirming that these structures are *bona fide* coiled-coils.

Our analysis of the interfaces between the coiled-coil α-helices indicates that the free energy of a human BicD2 with either asymmetric or homotypic registry would be expected to be similar. However, a human BicD2 with asymmetric registry would likely be somewhat more stable compared to BicD2 with homotypic registry, since additional non-covalent interactions are formed. This finding is further supported by the similar apparent melting temperatures of the *Dm* BicD-CTD and the human BicD2-CTD despite different registries in these structures. Notably, our CD spectra suggest that BicD2-CTD undergoes reversible structural changes at physiological temperatures (Figure 6C).

Together, our data support the hypothesis that human BicD2 exists in equilibrium between distinct registry-shifted conformers. As the auto-inhibited ground state, we propose BicD2 with an asymmetric registry, since this conformation is likely the most stable one (Figure 7). This conformation was also observed for *Dm* BicD that was crystallized in the absence of cargo (Liu et al., 2013), whereas homologs with homotypic registry were crystallized in the presence of cargo (Terawaki et al., 2015). Thus, cargo binding could induce a registry shift that converts the asymmetric coiled coil into a symmetric one (Figure 7). Such registry shift could activate auto-inhibited BicD2 for dynein recruitment upon binding of cargo. *In vivo* experiments are needed to validate this hypothesis. Our data suggests that binding of the cargo Rab^GTP^ is not affected by the F743I/R747C double mutation, but it remains to be established if a registry shift changes the selectivity of BicD2 towards other cargoes.

Notably, our data suggest a molecular mechanism for such a coiled-coil registry shift. In all conformations of BicD2 with homotypic registry, the side chains of residues F743 and F750 of both α-helices rotate outwards to be solvent exposed, which narrows the core of the coiled coil and allows the α-helices to be aligned in the homotypic registry. However, in conformations with asymmetric registry, the side chains of F743 and F750 are oriented towards the core of the coiled coil. As bulky aromatic residues are rare at core positions of coiled coils, residues F743, F750 and Y757 are accommodated by a large offset in their axial positions, resulting in the heterotypic registry. Of note, these orientations of F743 and F750 are directly observed in the simulation of the BicD2 R747C human disease variant that causes a spontaneous local registry shift, further confirming this mechanism.

Notably, in the classic *Drosophila* Bicaudal mutant, which results in a phenotype that includes double-abdomen flies, residue F684 (F743 in human BicD2) is mutated to isoleucine (Mohler and Wieschaus, 1986). The F684I mutation causes mRNA transport defects and does not alter cargo-binding. Instead, it leads to increased association of *Dm* BicD with dynein, thereby “hyperactivating” BicD2-dependent transport (Liu et al., 2013; Mohler and Wieschaus, 1986). Another residue linked to a registry shift is F750. A phenylalanine to isoleucine mutation of the homologous residue F691 in *Dm* BicD (Liu et al., 2013) causes a similar phenotype as the classic Bicaudal mutant. Thus, while residues F684 and F691 are located in the cargo-binding domain, they regulate recruitment of dynein in the NTD and likely play a key role in auto-inhibition. Our data suggest a potential mechanism of how the F684I mutation may lead to loss of autoinhibition of BicD2, since the homologous human residue F743 likely has a key role in facilitating coiled-coil registry shifts. In fact, in our MD simulations of the F743I variant, a spontaneous registry shift to a fully heterotypic registry was observed, further supporting the idea that a registry shift may be linked to BicD2-autoinhibiton.

**Figure 7.**
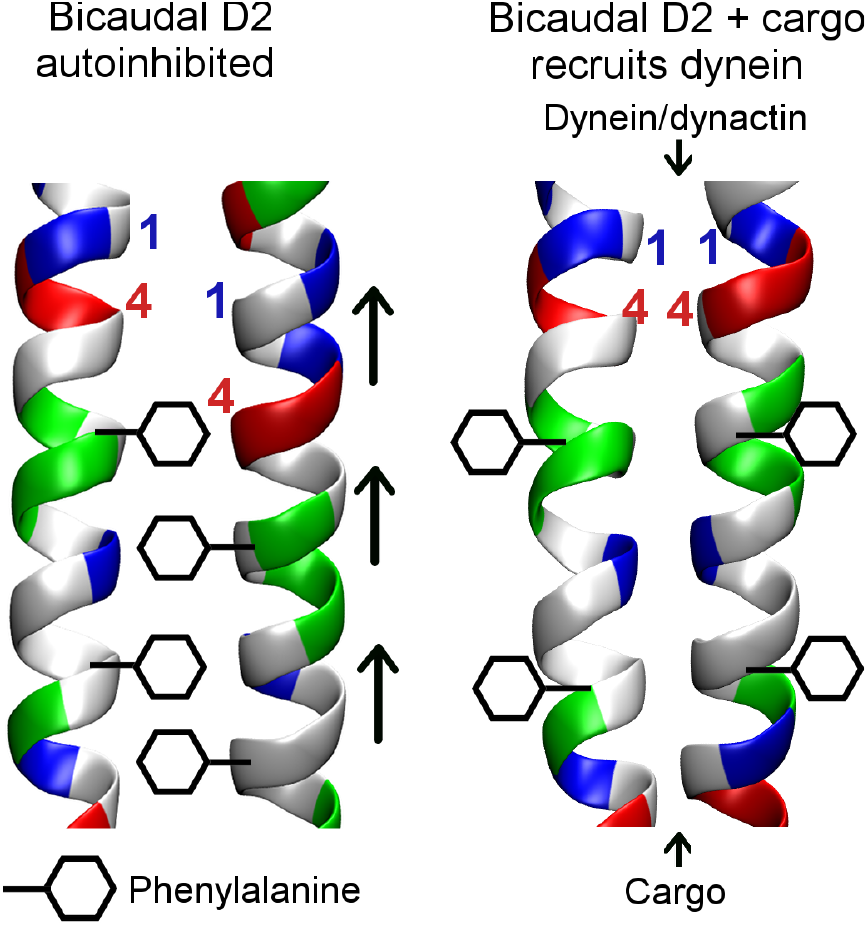
We propose BicD2 with an asymmetric registry as auto-inhibited ground state (left). Cargo binding could induce a registry shift that converts the asymmetric coiled coil into a symmetric one (right). These coiled coil registry shifts are likely facilitated by specific conformations of phenylalanine residues 743 and 750.

Coiled-coil registry shifts have also been reported for other cases (Carter et al., 2008; Choi et al., 2011; Croasdale et al., 2011; Gibbons et al., 2005; Macheboeuf et al., 2011; Snoberger et al., 2018; Xi et al., 2012). However, to our knowledge, MD simulations have only been used in one other study to model registry-shifted conformers of the coiled-coil stalk domain of dynein (Choi et al., 2011). During the power stroke, a registry shift in the stalk domain coordinates ATP hydrolysis with microtubule binding (Carter et al., 2008; Gibbons et al., 2005). In contrast to this former study, we directly observed a conversion of registry-shifted conformers in our simulations and we provide an in-depth analysis of the coiled-coil packing. Notably, all resulting structures that were analyzed featured good stereochemistry and characteristic knobs-into-holes interactions, confirming that *bona fide* coiled-coil structures with alternative registries can be formed. Notably, these structures are of comparable stability as the experimentally observed structures. These results suggest that the ability to undergo registry shifts may be an inherent property of coiled-coil structures, and likely more common than previously assumed.

Understanding how adaptors such as BicD2 modulate dynein motility is important as dynein facilitates a vast number of cellular transport events that are essential for chromosome segregation, neurotransmission, signaling, mRNA transport, and brain and muscle development. Our data support the notion that human BicD2 may undergo coiled-coil registry shifts, facilitated by specific conformations of residues F743 and F750. A registry shift upon cargo binding could propagate through the coiled-coil towards the N-terminal domain involved in dynein/dynactin binding in order to relieve autoinhibition of BicD2 and activate it for dynein recruitment. Of note, residue F743 is mutated to isoleucine in the variant that causes the Bicaudal phenotype in the *Drosophila* homolog, thereby linking a possible registry shift to auto-inhibition. Furthermore, we provide evidence that disease-causing mutations in coiled-coils may alter the equilibrium between registry-shifted conformers, which we propose as a general mechanism of pathogenesis in this class of proteins. The ability to undergo registry shifts may be an inherent and widespread property of coiled-coil structures. Thus, our results provide novel insights into an important dynamic property of coiled-coils, which are one of the most common tertiary structures.

## ACKNOWLEDGMENTS

This paper was supported by the following grants: National Institute of Health, National Institute of General Medical Sciences (NIH NIGMS) grant 1R15GM128119-01 awarded to S.R.S., the Chemistry Department at State University of New York at Binghamton and the Research Foundation of the State University of New York.

X-ray diffraction data was collected at beamline F1 at the MacCHESS facility of the Cornell High Energy Synchrotron Source, Ithaca, NY, which was supported by National Science Foundation (NSF) award DMR-1332208, and by NIH NIGMS award GM-103485. This work used the Extreme Science and Engineering Discovery Environment (XSEDE),(Towns et al., 2014) which is supported by NSF grant ACI-1548562. We thank A. Akhmanova (Utrecht University) for plasmids (Splinter et al., 2010), D. King (HHMI, UC Berkeley), for mass spectrometry analysis and S. Bane & B. Callahan (SUNY Binghamton), for access to equipment.

## AUTHOR CONTRIBUTIONS

C.R.N. and J.Y.L. performed research, analyzed data and made figures, C.R.N. designed research, E.W.D. analyzed data, K.M.L., H.C., and B.B.R. performed research. S.R.S and P.G. designed research, performed research, analyzed data, made figures and wrote the paper.

## DECLARATIONS OF INTERESTS

The authors declare no competing interests.

## ACCESSION NUMBERS

Coordinates and structure factors were deposited to the protein data bank (PDB ID 6OFP).

## SUPPLEMENTAL INFORMATION

Supplemental Information includes 4 figures and 4 tables, as well as supplementary methods and can be found with this article online.

## METHODS

### Structure determination of the human BicD2-CTD

Purified BicD2-CTD/Nup358 complex was crystallized in hanging drop setups at 20 °C. Crystals of the human BicD2-CTD (residues 715-804) were obtained in space group C2. For cryoprotection, crystals were briefly soaked in reservoir solution with addition of 30% glycerol and flash-frozen in liquid nitrogen.

X-ray diffraction images were collected at beamline F1, in the Macromolecular X-ray Science Facility at the Cornell High Energy Synchrotron Source, Ithaca, NY, from a single crystal. The structure was determined by molecular replacement in the PHENIX suite (Adams et al., 2010) with the structure of mouse BicD1 (Terawaki et al., 2015) as a search model, which was truncated at the N- and C-terminus. The structure was refined to 2.0 Å resolution, with an R_Free-_factor of 25.7%. The stereochemical quality of the model was assessed with MolProbity (Chen et al., 2010). The crystallographic statistics are summarized in Table S1.

### CD spectroscopy

CD spectroscopy of purified proteins was performed with a Jasco J-810 CD Spectrometer. After the buffer baseline subtraction, CD signals were normalized to protein concentration and converted to mean residue molar ellipticity [⊝]. Thermal unfolding profiles of proteins were recorded by CD spectroscopy at 222 nm. All proteins revealed a sigmoidal denaturation profile, characteristic of a two-state transition from helix to random coil, with a single melting temperature (T_M_). Therefore, for analysis of thermal unfolding profiles, a two-state denaturation model (α-helix to random coil) was used.

### MD simulations

Molecular dynamics simulations with implicit solvent were carried out using the CPU implementation of the PMEMD program in the AMBER16 package (Case et al., 2016). The solvent effects were mimicked by the GB-Neck2 implicit model (Mongan et al., 2007; Nguyen et al., 2013), where the surface area was computed using the linear combinations of pairwise overlaps (LCPO) model and the maximum distance between atom pairs that was considered when calculating the effective Born radii was 20.0 Å. The salt concentration was set at 0.15M. The ff14SBonlysc protein force field, mbondi3 intrinsic radii, and non-bonded cutoff distance of 50.0 Å were used in all calculations. The use of an implicit solvent model and a non-bonded cutoff distance of 50.0 Å were justified by comprehensive comparisons of the results to those from explicit solvent simulations as well as implicit solvent simulations with other values of the non-bonded cutoff distance.

To create an initial heterotypic BicD2-CTD structure, chain A in the experimental crystal structure of BicD2-CTD was manually pulled down by one helical turn relative to chain B. To create an initial structure of BicD2-CTD with asymmetric homotypic-heterotypic registry, the crystal structure of the CTD from *Dm* BicD (Liu et al., 2013) was mutated to match the sequence of human BicD2. For the R747C and F743I BicD2 mutants, initial structures were created by mutation of R747 in the experimental crystal structure and F743 in the equilibrated asymmetric registry structure, respectively. Further simulation procedures are provided in the Supplemental Information. MolProbity (Chen et al., 2010) structure validation, analysis of the knobs-into-holes interactions using SOCKET (Walshaw and Woolfson, 2001), and analysis of the dimer interface using PDBePISA (Krissinel and Henrick, 2007) were performed on the final structure from each simulation after subjecting it to an energy minimization.

