## Supplemental Information for "Role of coiled-coil registry shifts in activation of human Bicaudal D2 for dynein recruitment upon cargo-binding"

**The PDF file includes:**

| | | Page |
|---|---|---|
| 1. | Supplementary Methods | S2 |
| 2. | Figure S1 Knobs-into-holes interactions identified in the crystal structures | S7 |
| 3. | Figure S2 Structure of the human BicD2-CTD with heterotypic registry created *in silico* | S9 |
| 4. | Figure S3 Knobs-into-holes interactions identified in the simulation structures | S10 |
| 5. | Figure S4 Thermal unfolding curves of wild type BicD2-CTD and mutants | S11 |
| 6. | Table S1. Crystallographic statistics | S12 |
| 7. | Table S2. Dimer interfaces of BicD2-CTD homologs | S13 |
| 8. | Table S3. MolProbity structure validation reports for three structures of the BicD2-CTD with distinct coiled-coil registries | S14 |
| 9. | Table S4. Dimer interfaces for the structures of BicD2-CTD mutants | S15 |

### SUPPLEMENTARY INFORMATION TEXT

### SUPPLEMENTARY METHODS

#### Protein expression and purification

***Expression constructs:*** DNA sequences encoding human BicD2, *Drosophila* BicD, and human Nup358 were cloned into expression vectors as described (Cui et al., 2018; Noell et al., 2018). Plasmids encoding BicD2 and Nup358 fragments were provided by Anna Akhmanova (Utrecht University, the Netherlands) (Splinter et al., 2010). Expression constructs were generated by PCR from these templates and cloned into an expression vector by two restriction enzymes. Site directed mutagenesis was either performed as described (Cui et al., 2018) or performed by a commercial cloning service (Genscript).

***Protein expression and purification:*** All proteins were expressed in the *E. coli* Rosetta 2 BL21(DE3)pLysS strain at 37 °C as described (Cui et al., 2018). For BicD2-CTD/F743I, BicD2-CTD/F743I/R747C and the corresponding WT control, expression was performed in the LOBSTR BL21(DE3)-RIL strain. *Dm* BicD-CTD was expressed in the BL21(DE3)-RIL strain. Protein purification was performed using protocols as described (Cui et al., 2018; Noell et al., 2018; Solmaz et al., 2013; Solmaz et al., 2011), using the following strategies:

$His_6$-tagged BicD2-CTD and $His_6$-tagged BicD, as well as the BicD2-CTD variants were purified by Ni-NTA affinity chromatography, followed by protease cleavage of the $His_6$-tag (using thrombin or PreScission protease depending on the vector), and a second affinity chromatography step post-cleavage.

GST-tagged Nup358 was purified by glutathione affinity chromatography and eluted by PreScission protease cleavage of the tag. All proteins (with exception of the BicD2-CTD variants R747C/F743I and F743I, as well as the corresponding WT controls) were further purified by gel filtration chromatography as described (Cui et al., 2018; Noell et al., 2018).

Protein concentrations were determined by spectrophotometry with the peptide method and concentrated proteins were flash-frozen in liquid nitrogen as described (Cui et al.,

2018). Purified proteins were analyzed by SDS-PAGE, using 10% and 16% acrylamide gels, and stained by Coomassie Blue.

***GST-pulldown assay:*** Rab6$^{GTP}$ was purified by glutathione affinity chromatography as described (Noell et al., 2018), but not eluted. 1 mM GTP was added to the immobilized Rab6$^{GTP}$ on 0.5 ml of the resin. Purified BicD2-CTD WT or the F743I/R747C variant were added to the resin in four fold molar excess. The resin was incubated for 30 min while gently shaking and then washed with binding buffer. Elution was performed with 10 mM glutathione, 50 mM Tris pH 8.0, 150 mM NaCl, 1 mM DTT.

**Structure determination**

***Structure determination:*** X-ray intensities were processed and scaled using the programs XDS and XSCALE (Kabsch, 2010). The structure was determined by molecular replacement in the PHENIX suite (Adams et al., 2010) with the structure of mouse BicD1 (Terawaki et al., 2015) as a search model, which was truncated at the N- and C-terminus. An initial model was obtained from automatic model building in PHENIX and completed by manual model building in the program COOT (Adams et al., 2010; Emsley et al., 2010). The structure was refined to through iterative cycles of manual model building and refinement in PHENIX and COOT (Adams et al., 2010; Emsley et al., 2010). The structure was refined to 2.0 Å resolution, with an $R_{Free-}$factor of 25.7%.

***Structural analysis:*** Structures were compared by least-squares superimposition of the coordinates in COOT (Emsley et al., 2010). Dimer interfaces and knobs-into-holes interactions were analyzed by the web servers PISA and SOCKET, respectively (Krissinel and Henrick, 2007; Walshaw and Woolfson, 2001). For identification of knobs-into-hole interactions, a cutoff of 7.5 Å was used (helix extension 1 residue). Figures were created in the PyMOL Molecular Graphics System, Version 2.0 (Schrödinger, LLC) and VMD (Humphrey et al., 1996). Sequence alignments were performed with Clustal Omega (Sievers et al., 2011).

### CD Spectroscopy

***CD spectroscopy*** of purified proteins was performed with a Jasco J-810 CD Spectrometer equipped with a thermoelectric control device. Prior to analysis, purified proteins (at a concentration of 0.3 mg/ml) were dialyzed into 500 mM NaCl, 0.2 mM TCEP and 10 mM Tris, pH 8. For *Dm* BicD and the corresponding BicD2-CTD WT control, 150 mM NaCl was used. For melting curves in Figure 6H, a protein concentration of 0.5 mg/ml was used. The cuvette pathlength was 0.1 cm.

***CD wavelength spectra.*** CD spectra in the wavelength range of 200 to 250 nm were recorded using 1 nm bandwidth, 20 nm/min increments, 1 s equilibration time, 3 s accumulation time, at temperatures of 4°C or 90°C and data was averaged over three scans. Spectra were smoothened by a Savitsky-Golay algorithm in Origins Pro software (OriginLab Corp.) with a polynomial order of 3 and a smoothing window of 20 points. Multiple spectra for each protein were collected and representative data is shown.

***Thermal unfolding curves.*** Thermal unfolding data were recorded at 222 nm. The protein sample was heated in increments 1°C/min with 30 s equilibration time at each temperature, 1 nm bandwidth and 3 s accumulation time at each temperature. The apparent melting temperature ($T_M$) was determined from the peak of differential melting curves d[Θ222](T)/dT using Origins Pro software. Melting curves were also smoothed as described above. $T_M$ was averaged from three experiments, representative data are shown. Standard deviations were calculated. The fraction of folded peptide ($F_{folded}$) was calculated from the equation $F_{folded} = ([\Theta]-[\Theta]_{unfolded})/([\Theta]_{folded}-[\Theta]_{unfolded})$, where [Θ] is the observed molar ellipticity at any particular temperature and $[\Theta]_{unfolded}$ and $[\Theta]_{folded}$ are the molar ellipticities of the denatured (unfolded) and native (folded).

### Molecular dynamics simulations

***For the structure of wild-type homotypic BicD2-CTD***, hydrogens atoms were added to the starting crystal structure with the “reduce” functionality in AMBER (Case et al., 2016). Details of the implicit solvent model and the force field used are described in the main paper. 1000 steps of steepest descent geometry optimization followed by 2500 steps of conjugate gradient geometry optimization were carried out with a gradient tolerance of 0.02 kcal $mol^{-1}$ $Å^{-1}$. The system was then heated gradually to 300K and equilibrated for 1 ns prior to the production simulation. The SHAKE algorithm (Ryckaert et al., 1977) was used to constrain bonds involving hydrogen atoms. For temperature regulation, Langevin dynamics with a collision frequency of 1.0 $ps^{-1}$ was used. Subsequently, a production simulation at 300 K was performed for 50 ns with a timestep of 2 fs. Structure validation, and detailed analysis of the knobs-into-holes interactions and the dimer interface were performed on the final structure after subjecting it to an energy minimization.

***To create an initial heterotypic BicD2-CTD structure***, chain A in the crystal structure was manually pulled down by one helical turn relative to chain B. Hydrogen atoms were then added with the “reduce” functionality in AMBER (Case et al., 2016). Details of the implicit solvent model and the force field used are described in the main paper. 1000 steps of steepest descent geometry optimization followed by 6000 steps of conjugate gradient geometry optimization were carried out with a gradient tolerance of 0.02 kcal $mol^{-1}$ $Å^{-1}$. The system was then heated gradually to 300 K and equilibrated for 80 ns with harmonic restraints on the backbone atoms with a force constant of 10.0 kcal $mol^{-1}$ $Å^{-2}$. The SHAKE algorithm was used to constrain bonds involving hydrogen atoms. For temperature regulation, Langevin dynamics with a collision frequency of 1.0 $ps^{-1}$ was used. Subsequently, a production simulation with no restraints at 300 K was performed for 100 ns with a timestep of 2 fs. Structure validation, and detailed analysis of the knobs-into-holes interactions and the dimer interface were performed on the final structure after subjecting it to an energy minimization.

***To create an initial structure of BicD2-CTD with asymmetric homotypic-heterotypic registry***, the crystal structure of the CTD of *Dm* BicD (Liu et al., 2013) was mutated to match the sequence of human BicD2-CTD. Details of the implicit solvent model and the force field used are described in the main paper. 1000 steps of steepest descent geometry optimization followed by 200 steps of conjugate gradient geometry optimization were carried out with a gradient tolerance of 0.02 kcal $mol^{-1}$ $Å^{-1}$. The system was then heated gradually to 300K and equilibrated for 100 ns with harmonic restraints on the backbone atoms with a force constant of 10.0 kcal $mol^{-1}$ $Å^{-2}$. The SHAKE algorithm was used to constrain bonds involving hydrogen atoms. For temperature regulation, Langevin dynamics with a collision frequency of 1.0 $ps^{-1}$ was used. Subsequently, a production simulation with no restraints at 300 K was performed for 120 ns with a timestep of 2 fs. Structure validation, and detailed analysis of the knobs-into-holes interactions and the dimer interface were performed on the final structure after subjecting it to an energy minimization.

***For the R747C and F743I BicD2-CTD mutants***, after mutation of R747 in the experimental crystal structure and F743 in the equilibrated asymmetric registry structure, respectively, an equilibration procedure similar to that for the wild type homotypic BicD2-CTD was adopted. Subsequently, a production simulation at 300 K was performed for 100 ns for each mutant. Detailed analysis of the knobs-into-holes interactions and the dimer interfaces was performed on the final structures after subjecting them to energy minimization.

### SUPPLEMENTARY FIGURES AND TABLES

#### (A) Assignment of heptad repeats for BicD2-CTD

```
Chain A
Sequence GSHMYENEKAMVTETMMKLRNELKALKEDAATFSSLRAMFATRCDEYITQLDEMQRQLAAAEDEKKTLNSLLRMAIQQKLALTQRLELLEL
Register            defgabcdefgabcdefgabcdefgabcdefgabcdefgabcdefgabcdefgabcdefgabcdefgabcdefgabcd
Partner  ---------------Y--Y---Y--Y-----------------Y--Y---Y--Y---Y--Y---Y--Y---Y--Y---Y--Y---Y-----

Chain B
Sequence GSHMYENEKAMVTETMMKLRNELKALKEDAATFSSLRAMFATRCDEYITQLDEMQRQLAAAEDEKKTLNSLLRMAIQQKLALTQRLELLELD
Register            defgabcdefgabcdefgabcdefgabcdefgabcdefgabcdefgabcdefgabcdefgabcdefgabcdefgabcd
Partner  ---------------X--X---X--X---X--X------X------X---X--X---X--X---X--X---X--X---X--X---X------
```

#### (B) Assignment of heptad repeats for *Mm* BicD1-CTD

```
Chain A
GNKYENEKAXVTETXTKLRNELKALKEDAATFSSLRAXFATRCDEYVTQLDEXQRQLAAAEDEKKTLNTLLRXAIQQKLALTQRLEDLEFDHEQ
defgabcdefgabcdefgabcdefgabcdefgabcdefgabcdefgabcdefgabcdefgabcdefgabcdefgabcd
--------------Y--Y---Y--Y-----------------Y--Y---Y--Y---Y--Y---Y--Y---Y--Y------Y---Y---------

Chain B
GNKYENEKAXVTETXTKLRNELKALKEDAATFSSLRAXFATRCDEYVTQLDEXQRQLAAAEDEKKTLNTLLRXAIQQKLALTQRLEDLEFDHEQSRRSK
defgabcdefgabcdefgabcdefgabcdefgabcdefgabcdefgabcdefgabcdefgabcdefgabcdefgabcd
--------------X--X---X--X---X--X-------------X---X--X---X--X---X--X---X--X------X---X--X-----------
```

#### (C) Assignment of heptad repeats for *Dm* BicD-CTD

```
Chain A
VSDTMSKLRNELRLLKEDAATFSSLRAMFAARCEEYVTQVDDLNRQLEAAEEEKKTLNQLLRLAVQQKLALTQRLEEM
           abcdefgabcdefgabcdefgabcdefgabcdefgabcdefgabcdefgabcdefgabcdefga
-------Y---Y--Y----------Y-------------Y--Y---Y--Y---Y--Y---Y--Y------Y-------

Chain D
NEKIIVSDTMSKLRNELRLLKEDAATFSSLRAMFAARCEEYVTQVDDLNRQLEAAEEEKKTLNQLLRLAVQQKLALTQRLEEM
     abcdefgabcdefgabcdefgabcdefgabcdefgabcdefgdefgabcdefgabcdefgabcdefgabcdefga
!----X--X---X--X----------X------X-------------X---X--X---X--X---X---------X-------
```

#### (D) Sequence alignment of BicD2-CTD and *Mm* BicD1-CTD

```
 GSHMYENEKAMVTETMMKLRNELKALKEDAATFSSLRAMFATRCDEYITQLDEMQRQLA
GPGNKYENEKAMVTETMTKLRNELKALKEDAATFSSLRAMFATRCDEYVTQLDEMQRQLA
     ************ ******************************:***********

AAEDEKKTLNSLLRMAIQQKLALTQRLELLELDHE------
AAEDEKKTLNTLLRMAIQQKLALTQRLEDLEFDHEQSRRSK
**********:***************** **:***
```

#### (E) Sequence alignment of BicD2-CTD and *Dm* BicD-CTD

```
GSHMYENEKAMVTETMMKLRNELKALKEDAATFSSLRAMFATRCDEYITQLDEMQRQLAA
GSHMYENEKIIVSDTMSKLRNELRLLKEDAATFSSLRAMFAARCEEYVTQVDDLNRQLEA
********* :*::** ******: ****************:**:**:**:*:::*** *

AEDEKKTLNSLLRMAIQQKLALTQRLELLELDHE
AEEEKKTLNQLLRLAVQQKLALTQRLEEMEMDRE
**:******.***:*:*********** :*:*:*
```

**Figure S1A.** Related to Figure 2. Knobs-into-holes interactions were identified in the crystal structures (Liu et al., 2013; Terawaki et al., 2015) with the program SOCKET (Walshaw and Woolfson, 2001), and are denoted by ‘Y’ and ‘X’. Knob residues are located at ‘a’ and ‘d’ positions.

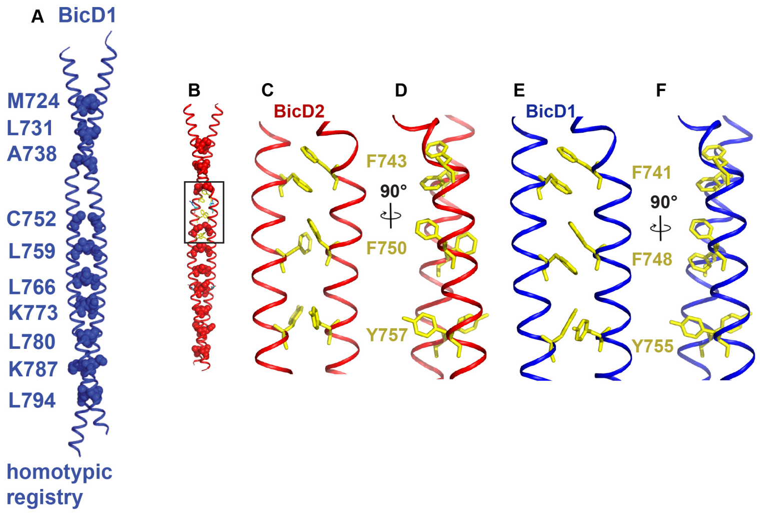


**Figure S1B.** Related to Figure 2. Comparison of the BicD2-CTD and BicD1-CTD (Terawaki et al., 2015) crystal structures. (A) Structure of mouse BicD1-CTD in cartoon representation. Residues in ‘a’ position of the heptad repeat are shown in spheres representation and labeled. (B) Structure of the human BicD2-CTD. Residues F743, F750 and Y757 are shown in stick representation (yellow). The boxed area is shown enlarged in (C) and rotated by 90° in (D). (E-F) The structure of the BicD1-CTD, homologous aromatic residues are highlighted in yellow and labeled. Note that the aromatic residues engage in distinct types of aromatic interactions in the structures of the homologs.

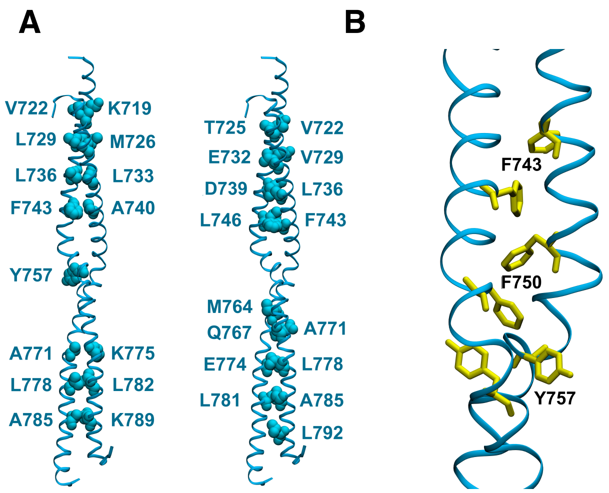


**Figure S2.** Related to Figure 4. The structure of the human BicD2-CTD with heterotypic registry created *in silico* is a *bona fide* coiled-coil. (A) Equilibrated and energy-minimized structure of the human BicD2-CTD with heterotypic registry at 300 K. Residues in 'a' position of the heptad repeat are shown as spheres in the left panel, and residues in 'd' position of the heptad repeat are shown as spheres in the right panel. The protein backbone is depicted in cartoon representation. (B) Orientation of key aromatic residues in the structure with the heterotypic registry: F743 and F750 are part of the dimer interface, while Y757 points outwards into solution.

#### (A) Assignment of heptad repeats for wild type BicD2-CTD, homotypic registry

```
Chain A
Sequence YENEKAMVTETMMKLRNELKALKEDAATFSSLRAMFATRCDEYITQLDEMQRQLAAAEDEKKTLNSLLRMAIQQKLALTQRLE
Register     abcdefgabcdefgabcdefgabcdefgabcdefgabcdefgabcdefgabcdefgabcdefgabcdefgabcdefga
Partner  -------Y---Y--Y---Y--Y---Y-------------Y------Y--Y---Y--Y---Y--Y---Y--Y------Y---Y-

Chain B
Sequence AMVTETMMKLRNELKALKEDAATFSSLRAMFATRCDEYITQLDEMQRQLAAAEDEKKTLNSLLRMAIQQKLALTQRLELLE
Register bcdefgabcdefgabcdefgabcdefgabcdefgabcdefgabcdefgabcdefgabcdefgabcdefgabcdefga
Partner  --X---X--X---X--X---X-------------X------X--X---X--X---X--X---X--X------X--------
```

#### (B) Assignment of heptad repeats for wild type BicD2-CTD, heterotypic registry

```
Chain A
Sequence MYENEKAMVTETMMKLRNELKALKEDAATFSSLRAMFATRCDEYITQLDEMQRQLAAAEDEKKTLNSLLRMAIQQKLALTQRLE
Register  defgabcdefgabcdefgabcdefgabcdefgabcdefgabcdefgabcdefdefgabcdefgabcdefgabcd
Partner  -----Y--Y---Y--Y---Y--Y---Y--Y--------------------Y--Y---Y--Y---Y--Y---Y------------

Chain B
Sequence MYENEKAMVTETMMKLRNELKALKEDAATFSSLRAMFATRCDEYITQLDEMQRQLAAAEDEKKTLNSLLRMAIQQKLALTQRLEL
Register     defgabcdefgabcdefgabcdefgabcdefgabcdefgabcdefgabcdefgdefgabcdefgabcdefgabcd
Partner  --------X--X---X--X---X--X---X--X----------XX------------X---X--X---X--X---X--X------
```

#### (C) Assignment of heptad repeats for wild type BicD2-CTD, asymmetric registry

```
Chain A
Sequence TMMKLRNELKALKEDAATFSSLRAMFATRCDEYITQLDEMQRQLAAAEDEKKTLNSLLRMAIQQKLALTQ
Register     defgabcdefgabcdefgabcdefgabcdefgabcdefgabcdefgabcdefgabcdefgabcd
Partner  ----Y---Y--Y---Y--Y------Y---Y--Y------Y---Y--Y---Y--Y---Y--Y---Y-----

Chain D
Sequence EKAMVTETMMKLRNELKALKEDAATFSSLRAMFATRCDEYITQLDEMQRQLAAAEDEKKTLNSLLRMAIQQKLALT
Register        defgabcdefgabcdefgabcdefgabcdefgabcdefgdefgabcdefgabcdefgabcdefgabcd
Partner  -----------X--X---X--X------X---X------X------X---X--X---X--X---X--X---X----
```

#### (D) Assignment of heptad repeats for R747C mutant of BicD2-CTD (homotypic registry). The mutation site is highlighted in blue. New or missing knobs compared to wild type BicD2-CTD, homotypic registry, are highlighted in green.

```
Chain A
Sequence XMYENEKAMVTETMMKLRNELKALKEDAATFSSLCAMFATRCDEYITQLDEMQRQLAAAEDEKKTLNSLLRMAIQQKLALTQRLEL
Register       abcdefgabcdefgabcdefgabcdefgabcdefgabcdefgabcdefgabcdefgabcdefgabcdefgabcdefga
Partner  ---------Y---Y--Y---Y--Y---Y------Y------Y------Y--Y---Y--Y---Y--Y---Y---------Y------

Chain B
Sequence YENEKAMVTETMMKLRNELKALKEDAATFSSLCAMFATRCDEYITQLDEMQRQLAAAEDEKKTLNSLLRMAIQQKLALTQRLEL
Register     abcdefgabcdefgabcdefgabcdefgabcdefgabcdefgabcdefgabcdefgabcdefgabcdefgabcdefga
Partner  -------X---X--X---X--X---X------X------X------X--X---X--X---X--X---X--X------X------
```

#### (E) Assignment of heptad repeats for F743I mutant of BicD2-CTD (heterotypic registry). The mutation site is highlighted in blue. New or missing knobs compared to wild type BicD2-CTD, heterotypic registry, are highlighted in green.

```
Chain A
Sequence MVTETMMKLRNELKALKEDAATISSLRAMFATRCDEYITQLDEMQRQLAAAEDEKKTLNSLLRMAIQQKLALTQRLELLE
Register  defgabcdefgabcdefgabcdefgabcdefgabcdefgabcdefgabcdefgabcdefgabcdefgabcdefgabcd
Partner  -----Y--Y---Y--Y------Y---Y------Y------Y--Y---Y--Y---Y--Y---Y--Y---Y--Y---Y--Y-

Chain D
Sequence EKAMVTETMMKLRNELKALKEDAATISSLRAMFATRCDEYITQLDEMQRQLAAAEDEKKTLNSLLRMAIQQKLALTQRLELL
Register defgabcdefgabcdefgabcdefgabcdefgabcdefgabcdefgabcdefgabcdefgabcdefgabcdefgabcd
Partner  ----X--X---X--X---X------X--X---X------X------X--X---X--X---X--X---X--X---X-------
```

**Figure S3.** Related to Figure 4 and Figure 5. Knobs-into-holes interactions were identified in the equilibrated and energy-minimized simulation structures with the program SOCKET using a distance cutoff of 7.5 Å and helix extension of 1 residue (Walshaw and Woolfson, 2001).

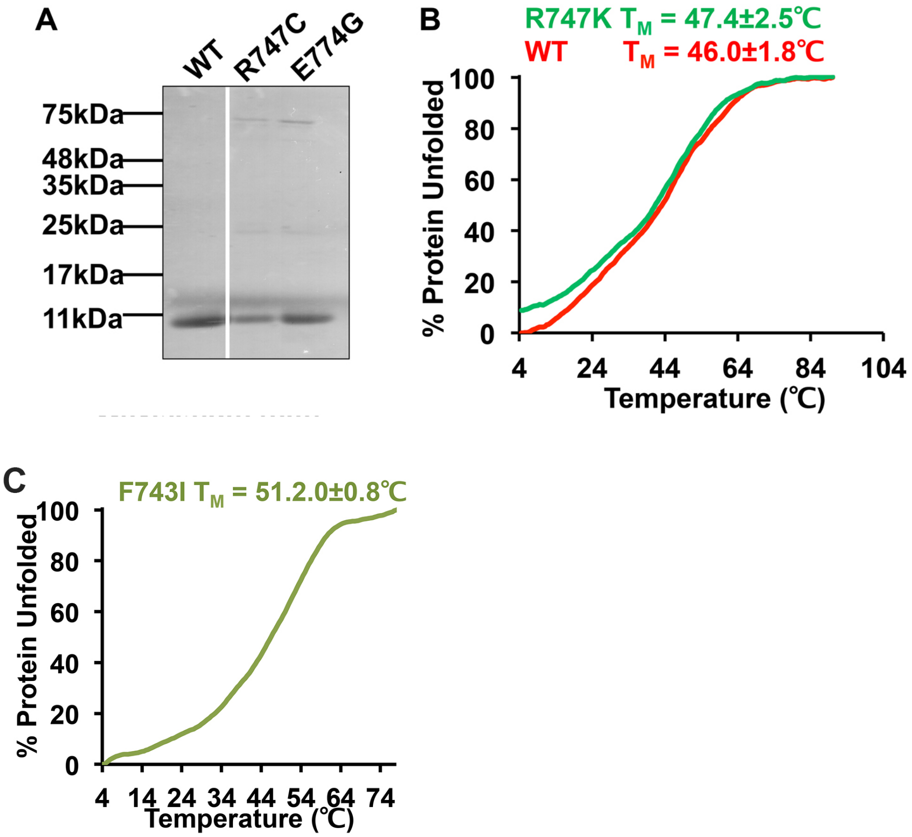


**Figure S4.** Related to Figure 6. (A) BicD2-CTD and the variants R747C and E774G were purified by Ni-affinity chromatography using His-Select affinity gel, and analyzed by SDS-PAGE. The molar masses of standard proteins are indicated. (B) Thermal unfolding curves of BicD2-CTD (red), and its variant R747K (green) monitored by CD spectroscopy at 222 nm. Unfolding is monitored by the increase of the molar ellipticity [Θ] at 222 nm as a function of temperature. 0% and 100% Protein unfolded represents the minimum and maximum value of Θ for wild-type BicD2-CTD, respectively. Apparent melting temperatures $T_M$ and standard deviations are shown. (C) Thermal unfolding curve of BicD2-CTD/F743I.

**Table S1. Related to Figure 1.**
**Crystallographic statistics**

| | |
|---|---|
| **Data collection statistics** | |
| Space group | *C*2 |
| Unit cell parameters a, b, c | 73.7, 37.0, 78.7 |
| Unit cell parameters $\alpha$, $\beta$, $\gamma$ | 90°, 107.4°, 90° |
| Wavelength (Å) | 0.98 |
| Resolution range (Å) | 40-2.0 (2.02-2.0)* |
| Rsym (%) | 7.0 (34.2)* |
| $I/\sigma(I)$ | 14.9 (2.0)* |
| Completeness (%) | 98.0 (85.0)* |
| Reflections meas./unique | 84722/25927 |
| Redundancy | 3.3 (2.2)* |
| **Refinement statistics** | |
| $R_{free}/R_{work}$ (%) | 25.7/22.6 |
| Reflections (work/test sets) | 11800/1313 |
| R.m.s.d. bonds (Å)/angles | 0.003/0.67 |
| **Ramachandran plot analysis** | |
| Favored | 100% |
| Allowed, outliers | 0% |

*Highest resolution shell in parenthesis. Meas: measured.

### Table S2. Related to Figure 2. Dimer interfaces of BicD2-CTD homologs

| ***Hs* BicD2** | | | ***Mm* BicD1** (PDB ID 4YTD) | | | ***Dm* BicD** (PDB ID 4BL6) | | |
|---|---|---|---|---|---|---|---|---|
| **Chain A** | **Chain B** | **Dist (Å)** | **Chain A** | **Chain B** | **Dist (Å)** | **Chain A** | **Chain D** | **Dist (Å)** |
| **Hydrogen bonds** | | | **Hydrogen bonds** | | | **Hydrogen bonds** | | |
| CYS 754 SG | CYS 754 SG | 3.9 | GLU 716 OE1 | TYR 713 OH | 2.8 | TYR 698 OH | CYS 695 SG | 3.2 |
| GLU 774 OE1 | LYS 775 NZ | 3.0 | GLU 772 OE1 | LYS 773 NZ | 2.6 | GLN 701 OE1 | TYR 698 OH | 3.0 |
| GLN 788 OE1 | LYS 789 NZ | 3.0 | | | | GLU 715 OE1 | LYS 716 NZ | 2.5 |
| LYS 775 NZ | GLU 774 OE1 | 2.3 | | | | ARG 688 NH1 | ASP 680 OD1 | 2.8 |
| | | | | | | ARG 688 NH2 | THR 683 OG1 | 2.8 |
| **Salt bridges** | | | **Salt bridges** | | | **Salt bridges** | | |
| GLU 732 OE2 | LYS 737 NZ | 3.4 | GLU 730 OE2 | LYS 735 NZ | 3.0 | GLU 715 OE1 | LYS 716 NZ | 2.5 |
| GLU 774 OE1 | LYS 775 NZ | 3.0 | GLU 772 OE1 | LYS 773 NZ | 2.6 | GLU 715 OE2 | LYS 716 NZ | 3.9 |
| LYS 737 NZ | GLU 732 OE1 | 3.1 | LYS 735 NZ | GLU 730 OE2 | 3.3 | ARG 688 NH1 | ASP 680 OD1 | 2.8 |
| LYS 775 NZ | GLU 774 OE1 | 2.3 | LYS 773 NZ | GLU 772 OE1 | 3.4 | ARG 688 NH2 | ASP 680 OD1 | 3.1 |
| | | | | | | LYS 716 NZ | GLU 715 OE2 | 2.9 |
| **Interface area** | | | **Interface area** | | | **Interface area** | | |
| 2161 Å$^2$ | | | 2176 Å$^2$ | | | 1764 Å$^2$ | | |
| **Interface residues** | | | **Interface residues** | | | **Interface residues** | | |
| 715, 718, 719, 722, 725, 726, 729, 730, 732, 733, 736, 737, 739, 740, 742, 743, 746, 747, 750, 751, 753, 754, 757, 758, 760, 761, 764, 765, 767, 768, 771, 772, 774, 775, 778, 779, 779, 792, 793, 795, 796, 799, 790 | | | 713, 716, 717, 720, 723, 724, 727, 728, 730, 731, 734, 735, 737, 738, 740, 741, 744, 745, 748, 751, 752, 755, 756, 758, 759, 762, 763, 765, 766, 769, 770, 772, 773, 776, 777, 779, 780, 783, 784, 786, 787, 790, 791, 793, 794, 797, 798, 801 | | | 659, 662, 663, 664, 666, 667, 669, 670, 671, 673, 674, 677, 678, 680, 681, 683, 684, 687, 688, 690, 691, 692, 694, 695, 698, 699, 701, 702, 705, 706, 708, 709, 712, 713, 715, 716, 719, 720, 722, 723, 726, 727, 729, 730, 733, 734, 736, 737, 740 | | |

Standard nomenclature from the structural coordinates is used for chain ID, residue names and atom names. Dist: Distance. Structure references:(Liu et al., 2013; Terawaki et al., 2015).

**Table S3. Related to Figure 4. MolProbity structure validation reports (Chen et al., 2010) for three possible conformations of the wild type BicD2-CTD coiled-coil, namely, homotypic registry, heterotypic registry and asymmetric registry.[a]**

| | BicD2-CTD, homotypic | BicD2-CTD, heterotypic | BicD2-CTD, asymmetric | Goal |
|---|---|---|---|---|
| **Clashscore, all atoms** | 0 | 0 | 0 | Clashscore is the number of serious steric overlaps (> 0.4 Å) per 1000 atoms |
| **Poor rotamers** | 2.55% | 1.27% | 1.39% | Goal: <0.3% of all rotamers |
| **Favored rotamers** | 96.18% | 96.18% | 95.83% | Goal: >98% of all rotamers |
| **Ramachandran favored** | 99.42% | 99.42% | 98.15% | Goal: >98% of all $\phi$, $\psi$ torsions |
| **$C_\beta$ deviations > 0.25 Å** | 0.00% | 0.00% | 0.60% | Goal: 0 |
| **Bad bonds** | 0.00% | 0.00% | 0.00% | Goal: 0% of all bonds |
| **Bad angles** | 0.41% | 0.51% | 0.39 % | Goal: <0.1% of all angles |

[a] Structures were energy-minimized post equilibration at 300 K. Poor/favored rotamers, favored Ramachandran dihedrals, $C_\beta$ deviations, and bad bonds/angles are determined as defined in Ref. (Chen et al., 2010).

**Table S4. Related to Figure 5 and Table 1. Dimer interfaces for the structures of wild-type BicD2-CTD with heterotypic registry and the variants BicD2-CTD/R747C and F743I.[a]**

| BicD2-CTD, heterotypic | | | BicD2-CTD/R747C | | | BicD2-CTD/F743I | | |
|---|---|---|---|---|---|---|---|---|
| Chain A | Chain B | Dist. (Å) | Chain A | Chain B | Dist. (Å) | Chain A | Chain B | Dist. (Å) |
| **Hydrogen bonds** | | | **Hydrogen bonds** | | | **Hydrogen bonds** | | |
| LYS 719 HZ3 | GLU 718 OE1 | 1.77 | LYS 719 HZ1 | GLU 718 OE2 | 1.79 | LYS 737 HZ1 | GLU 732 OE1 | 1.85 |
| ARG 730 HE | GLU 732 OE1 | 1.88 | LYS 775 HZ1 | GLU 774 OE1 | 1.85 | ARG 747 HH11 | ASP 739 OD1 | 1.82 |
| ARG 730 HH21 | GLU 732 OE1 | 2.00 | LYS 789 HZ2 | GLN 788 OE1 | 1.82 | LYS 775 HZ2 | GLN 767 OE1 | 1.93 |
| ARG 747 HE | MET 749 SD | 2.45 | GLU 718 OE1 | LYS 719 HZ1 | 1.78 | ASN 779 HD22 | GLU 775 OE1 | 1.81 |
| GLN 767 HE21 | GLU 772 OE2 | 1.96 | | | | LYS 789 HZ3 | MET 784 SD | 2.31 |
| LYS 775 HZ1 | GLU 774 OE1 | 1.81 | | | | GLU 772 OE2 | GLN 767 HE22 | 1.84 |
| GLU 774 OE2 | GLU 775 HZ2 | 1.78 | | | | GLU 774 OE1 | LYS 775 HZ3 | 1.82 |
| | | | | | | GLN 788 OE1 | LYS 789 HZ2 | 1.99 |
| | | | | | | GLU 800 OE1 | ARG 795 HH11 | 1.90 |
| **Salt bridges** | | | **Salt bridges** | | | **Salt bridges** | | |
| LYS 719 NZ | GLU 718 OE2 | 2.76 | LYS 719 NZ | GLU 718 OE2 | 2.79 | LYS 737 NZ | GLU 732 OE1 | 2.82 |
| ARG 730 NE | GLU 732 OE1 | 2.84 | LYS 775 NZ | GLU 774 OE1 | 2.78 | ARG 747 NE | ASP 739 OD1 | 3.98 |
| ARG 730 NH2 | GLU 732 OE1 | 2.91 | GLU 718 OE1 | LYS 719 NZ | 2.80 | ARG 747 NH1 | ASP 739 OD1 | 2.81 |
| LYS 775 NZ | GLU 774 OE1 | 2.80 | | | | GLU 774 NH1 | LYS 775 NZ | 2.81 |
| GLU 774 OE2 | LYS 775 NE | 2.78 | | | | GLU 800 OE1 | ARG 795 NH1 | 2.84 |
| **Interface Area (Å$^2$)** | | | **Interface Area (Å$^2$)** | | | **Interface Area (Å$^2$)** | | |
| 2339 | | | 2319 | | | 2251 | | |
| **Interface Residues** | | | **Interface Residues** | | | **Interface Residues** | | |
| 714, 715, 716, 718, 719, 721, 722, 723, 725, 726, 728, 729, 730, 732, 733, 735, 736, 737, 739, 740, 742, 743, 744, 746, 747, 749, 750, 751, 753, 754, 757, 758, 760, 761, 763, 764, 767, 768, 770, 771, 772, 774, 775, 777, 778, 779, 781, 782, 784, 785, 786, 788, 789, 791, 792, 793, 795, 796, 799, 801 | | | 714, 715, 718, 719, 721, 722, 723, 725, 726, 729, 730, 732, 733, 736, 737, 739, 740, 743, 744, 746, 747, 750, 751, 753, 754, 755, 757, 758, 760, 761, 764, 765, 767, 768, 771, 772, 774, 775, 778, 779, 781, 782, 784, 785, 786, 788, 789, 792, 793, 795, 796, 799 | | | 717, 718, 719, 721, 722, 723, 725, 726, 728, 729, 730, 732, 733, 736, 737, 739, 740, 742, 743, 744, 746, 747, 749, 750, 751, 753, 754, 756, 757, 758, 760, 761, 763, 764, 765, 767, 768, 770, 771, 772, 774, 775, 777, 778, 779, 781, 782, 784, 785, 786, 788, 789, 791, 792, 793, 795, 796, 798, 799, 800, 801 | | |

[a] Structures were energy-minimized post equilibration at 300 K. The F743I mutant has 9 and 8 fewer residues in chain A and B, respectively, compared to the R747C mutant and the wild-type structure with heterotypic registry. The F743I simulation used an initial structure with asymmetric registry while the R747C simulation used an initial structure with homotypic registry.
